# Alamandine/MrgD Pathway Modulates Gut-Bone Marrow Axis in Aging

**DOI:** 10.64898/2026.02.12.705187

**Authors:** Kishore Chittimalli, Henryata E Rozario, Victor Martinez, Zachary L McAdams, Stephen A Adkins, Aaron C Ericsson, Yagna PR Jarajapu

## Abstract

Aging is associated with colon epithelial barrier integrity and upregulation of myelopoiesis in the bone marrow (BM). Alamandine (Ala) and MrgD are novel members of the renin angiotensin system (RAS). This study tested the hypothesis that Ala restores the colon epithelial barrier integrity in aging via modulating gut-BM axis.

Mice of age 2-3 (Young) or 22-24 months (Old) were treated with saline or Ala by using Osmotic pumps. The intestinal permeability was evaluated by using FITC-dextran. Lgr5^+^Olfm4^+^ intestinal stem cells (ISCs), Wnt3a and β-catenin were evaluated by immunohistochemistry or western blotting. Fecal microbiome was analyzed by 16S rRNA sequencing. Monocyte-macrophages were characterized by flow cytometry. Cecal or serum bacterial metabolites were analyzed. The pro-myelopoietic potential of cecal supernatants (CS) was tested in the Young-BM cells.

MrgD was expressed in ISCs, which was decreased in the Old. Increased intestinal permeability in aging was reversed by Ala. In the colon organoids, Ala increased Wnt3a levels that were antagonized by the NF449, SQ22536 or 666-15. Ala restored phospho-CREB and active β-catenin levels that were decreased in the Old colon-organoids. Ala increased the richness and β-diversity of the aging microbiome and decreased *Bacillota*/*Bacteroidota*. Ala decreased the CD80^+^ and increased CX3CR^+^ cells in the Old colons. Old-CS induced myelopoiesis in vitro in BM cells with higher number of monocytes and pro-inflammatory macrophages which was not observed in the CS derived from Ala-treated Old mice.

Ala is a promising pharmacological agent for reversing the leaky gut of aging by restoring homeostasis in the gut-BM axis.

## Introduction

Aging is characterized by a decline in the multiorgan functions and an increased risk for the development of localized and systemic pathologies. Chronic low-grade systemic inflammation in aging, aka inflammaging, is manifested with local and systemic pathological phenotypes. Inflammaging largely originates from the myelopoietic bias in the bone marrow (BM) hematopoiesis with increased number of circulating monocytes and inflammatory cytokines and infiltration of peripheral tissues with pro-inflammatory macrophages (Mϕs).(1, 2) Monocyte-macrophages participate in physiological functions such as resolving inflammation (3) however a persistent increase in the circulating monocytes promotes tissue infiltration, resulting in systemic and local inflammation.

The disrupted colon epithelial barrier integrity, leaky gut, actively contributes to inflammaging. Intestine is a stem cell-based organ and the intestinal stem cells (ISCs) largely maintain the epithelial barrier integrity by regenerating multiple cell types including epithelial, goblet and Paneth cells as well as the molecular components of tight junctions, claudin 1, occludin and zonulin. (4) Proliferation, differentiation and transcription of junctional proteins by ISCs is dependent on wingless-related integration site (Wnt)/β-catenin signaling.(5–7) Aging is associated with decreased density of active ISCs due to downregulation of Wnt signaling leading to structural disruption of epithelial layer and disintegrity of tight junction proteins.(8) Reactivation of Wnt3a signaling restored the function of both murine and human ISCs and reversed the aging phenotype.(8) Gut microbiome has now emerged as a strong determinant of multiple physiological processes and overall maintenance of the host health.

However, the architecture of the gut microbiota undergoes progressive drift towards decreased ecological diversity aka microbial dysbiosis. Age-related changes in the microbiota promote disruption of the intestinal barrier resulting in the leakage of pro-inflammatory bacterial metabolites or membrane proteins such as lipopolysaccharide (LPS) that are TLR ligands promote inflammation in the peripheral tissues and induce myeloid skewing in the BM stem/progenitor cells as reported in humans and mice.(9, 10) Therefore, gut microbial dysbiosis in aging largely contributes to inflammaging.(11)

There is evidence for a bidirectional interaction of gut microbiome and BM cell populations via soluble metabolites derived from gut bacteria and paracrine factors released by BM cells. BM-derived monocytes migrate and infiltrate the colon wall and create local inflammation by differentiating into the pro-inflammatory Mϕs that promote gut barrier disintegrity and microbial dysbiosis.(12, 13) Fecal microbiome transfer reversed the changes induced by antibiotic in the BM hematopoiesis induced by antibiotics.(14) The differentiation potential of stem/progenitor cells was restored in the germ-free mice by recolonization of microbiota or by co-housing with conventionally raised mice(15). In support of this, bacterial metabolites were shown to modulate BM-hematopoiesis.(16, 17)

Renin-angiotensin system (RAS), a complex network of enzymes, small peptides and receptors, is yet expanding with novel peptide members and receptors. The two axes of RAS with opposing functions, the classical and the protective axes, are represented by angiotensin-converting enzyme/Angiotensin II/AT1 receptor and ACE2/Ang-(1–7)/Mas receptor (MasR), respectively.(18) Alamandine (Ala) is a relatively new peptide member of RAS, generated by either decarboxylation of the aspartic acid residue of Ang-(1–7) by an unidentified enzyme or by the enzymatic effect of ACE2 on Ang A.(19, 20)

Pharmacological profile of Ala in the cardiovascular tissues appears to be similar to that of Ang-(1–7) however mediated by a distinct receptor, Mas-related G-protein coupled receptor, member D (MrgprD or MrgD) aka Takeda G protein-coupled receptor 7 (TGR7).(19) Functional significance of MrgD was first reported in the dorsal root ganglion neurons that respond to pain stimuli and therefore proposed to be a novel target for the treatment of neuropathy.(21, 22) Later studies showed evidence for the expression of this receptor in several nonneuronal tissues. The report by Lautner et al (19) increased our understanding of Ala/MrgD pharmacology and the evidence for the protective role of Ala/MrgD pathway in cardiovascular diseases started accumulating.(23) Very recent study reported the beneficial effects of this pathway in rat and mouse models of pulmonary arterial hypertension (PAH) while MrgD-deficient mice showed exacerbated pathological phenotype when exposed to PAH-inducing insults.(24) To date, studies reported coupling of MrgD to different signaling pathways in different cell types based on the limited number of agonists that were characterized including β-alanine, Ala and a nonprotein amino acid L-β-aminoisobutyric acid (L-BAIBA).(19, 22, 25, 26) Gαi/0, Gαq/11 or Gαs -coupling was reported in different cell types largely based on the pharmacological inhibitors of signaling molecules.(22, 25, 27, 28)

We have previously shown experimental evidence for protective effects of Ang-(1–7) in the leaky gut of aging or diabetes largely by restoring the regenerative functions of ISCs via MasR/Wnt3 pathway.(10, 29) Pharmacology of Ala in the colon is unknown. This study aimed at unraveling the protective role of Ala/MrgD pathway on the gut-BM axis in aging. We have carried out *in vivo* pharmacology in mice and *in vitro*/*ex vivo* pharmacology in the cultured colon-organoids to evaluate the protective functions of Ala on the colon permeability and to delineate the cellular signaling of Ala/MrgD, respectively. Myelopoiesis and Mϕ polarization assays were carried out in the BM cells. Mechanistic interaction of gut metabolites and BM myelopoiesis was tested by using cecal supernatants on the myelopoietic functions of BM cells.

## Materials and Methods

### Animal model

All animal experiments were carried out at North Dakota State University and upon approval by the Institutional Animal Care and Use (IACUC) Committee. C57Bl/6 male and female mice were obtained from the National Institute of Aging (NIA) Rodent facility at Charles River Laboratory. Mice were housed in a conventional facility on a light-dark cycle of 12 hours provided with standard rodent chow and water *ad libitum*. Mice were of age of 3-4 (Young) or 22-24 (Old) months at the time experimentation and were housed in our facility for a minimum of three months in the case of Young or seven months in the case of Old prior to assigning to experimental groups.

### Pharmacological treatments

Treatment with saline, Ala or D-Pro^7^-Ang-(1–7) (Bachem and MedChem Express) was accomplished by using an osmotic pump (Alzet) implanted subcutaneously under isoflurane anesthesia. Osmotic pumps were prepared to release Ala or D-Pro^7^-Ang-(1–7) at a perfusion rate of 1 µg/kg/min for four weeks, as described before.(30–33) Where applicable, D-Pro^7^-Ang-(1–7) was administered along with Ala by using another osmotic pump. At the end of the treatment protocol, mice were euthanized by thoracotomy and cardiac puncture under isoflurane anesthesia.

### *In vivo* intestinal permeability

FITC-conjugated Dextran was used for the assessment of intestinal permeability. Fluorescein isothiocyanate (FITC)-dextran (Sigma Aldrich) in saline (400 mg/kg body weight, 125 mg/mL) was administered by oral gavage and blood samples were collected after 1, 2.5 and 4 hours and samples were processed as described before for spectrophotometric analysis of FITC-dextran.(10, 29)

### *Ex vivo* culture of colonic organoids

The *ex vivo* culture of organoids from mouse colons was carried out as described by Liu et al (34) by using Intesticult (StemCell Technologies). Colon crypts were collected from the dissociated colon fragments and mixed with Matrigel (Millipore Sigma) to plate in a 24 well plate. Where applicable, organoids were treated with Ala or Ang-(1–7) with or without A779 or D-Pro7-Ang-(1–7) at different concentrations.

### Western blotting

Western blotting was carried out by using standard laboratory procedures as described before Chittimalli 23 by using protein from the colon tissue lysates for electrophoresis using 4-20% SDS-PAGE gradient gels (Bio-rad). Proteins were transferred to PVDF membranes and prepared for treatment with primary and secondary antibodies. The protein bands were detected by using enhanced chemiluminiscence substrate (Thermo-Scientific) and the band intensities were quantified by using ImageJ software.(35) Sources and concentrations of different antibodies are listed in the supplementary Table S1.

### Histochemistry and immunocytochemistry

The intestinal barrier integrity was evaluated by Hematoxylin & Eosin (H&E) staining. Colon tissues were isolated and fixed in 4% Paraformaldehyde solution for 24 hours at room temperature, which were then embedded in paraffin blocks to obtain sections of five-micron thickness. The sections were deparaffinized using xylene then hydrated and stained using Harris Hematoxylin Solution (Electron Microscopy Sciences) - Eosin Y Phloxine B Solution (Electron Microscopy Sciences). Subsequently, the sections were mounted by using Permount the mounting medium, Permount (Electron Microscopy Sciences) and were imaged by using a fluorescence microscope (Leica).

The expression of claudin 1 or occludin and the presence of ISCs and inflammatory cells were characterized by immunocytochemistry. The deparaffinized sections were subjected to epitope retrieval process, which involved steaming in sodium citrate buffer pH 6.0. Then, normal goat serum was used to avoid non-specific binding and incubated with primary antibodies overnight at 4°C. Sections were washed with TBST and incubated with secondary antibodies for two hours at room temperature. Sections were washed with water and counterstained with DAPI and imaged by Zeiss Fluorescence microscope with 20× magnification. Negative controls were prepared by excluding primary antibodies. Sources and concentrations of different antibodies used for immunohistochemistry are listed in the supplementary Table S1.

### Biochemical analysis

Plasma LPS-binding protein (LBP) and Trimethyl amine N-oxide (TMAO) levels were analyzed by using an ELISA kits from Abcam and BioHippo, respectively.

### Microbiome analysis

Fecal sample collection and processing of samples for bacterial DNA extraction were carried out as described before.(10) Bacterial 16S rRNA amplicons were constructed by PCR amplification of the V4 hypervariable region 16S rRNA gene with universal primers (U515F/806R), flanked by Illumina standard adapter sequences. Amplified products were purified by Axygen AxyPrep MagPCR Clean-up beads and the final amplicon pool was evaluated by using the Advanced Analytical Fragment Analyzer automated electrophoresis system, quantified with the Qubit fluorometer by using the quant-iT BR dsDNA reagent kit, and prepared according to Illumina’s standard protocol for sequencing by MiSeq.

The DNA sequences were assembled and annotated at the MU Bioinformatics and Analytics Core facility. Primers were designed to match the 5’ ends of the forward and reverse reads. Cutadapt (36) (version 2.6; https://github.com/marcelm/cutadapt) was used to remove the primer from the 5’ end of the forward read. If found, the reverse complement of the primer to the reverse read was then removed from the forward read as were all bases downstream. Thus, a forward read could be trimmed at both ends if the insert was shorter than the amplicon length. The same approach was used on the reverse read, but with the primers in the opposite roles. Read pairs were rejected if one read or the other did not match a 5’ primer, and an error-rate of 0.1 was allowed. Two passes were made over each read to ensure removal of the second primer. A minimal overlap of three bp with the 3’ end of the primer sequence was required for removal. The QIIME2 (37) DADA2 (38) plugin (version 1.10.0) was used to denoise, de-replicate, and count ASVs (amplicon sequence variants), incorporating the following parameters: 1) forward and reverse reads were truncated to 150 bases, 2) forward and reverse reads with number of expected errors higher than 2.0 were discarded, and 3) Chimeras were detected using the “consensus” method and removed. R version 3.5.1 and Biom version 2.1.7 were used in QIIME2. Taxonomies were assigned to final sequences using the Silva.v132 (39) database, by using the classify-sklearn procedure.

Alpha and beta diversity metrics were determined by using R v4.2.2 (https://www.r-project.org). (40) By using the *microbiome* and *vegan* libraries, Chao1 richness and Shannon diversity indices were determined.(41, 42) Beta diversity with weighted Bray-Curtis distances was determined by using the *vegan* library. Principal coordinate analysis (PCoA) was performed by using the *ape* library.(43) Differences in the beta diversity were determined by permutational analysis of variance (PERMANOVA) using the *vegan* and *EcolUtils* (44) libraries with 9,999 permutations. The false discovery rate inherent in the differential abundance testing was controlled by using the method of Benjamini and Hochberg.(45) Spearman correlations, significance testing, and Benjamini-Hochberg corrections were performed using base R *stats* library.(40)

### Metabolomics Profiling

Serum and cecal contents were analyzed for bacterial metabolites and bile acid metabolites by Metabolomics core facility at the University of Iowa by using deuterated standard based LC-MS methodology. LC-MS data were processed by TraceFinder 4.1 software (Thermo Scientific), and metabolites were identified based on the standard-confirmed, inhouse library. Analyte signal was corrected by normalizing to the deuterated analyte signal and the with blank correction.

### Inflammatory Cytokine Microarray

Colon tissue lysates were used to analyze cytokines by using Raybiotech Mouse Inflammation Array Q1 (QAM-INFL-1). Protein arrays were prepared according to the manufacturer’s instructions and the fluorescent signals from the arrays were recorded by using Innopsys Innoscan 710 at the Raybiotech service center. Sample concentration in pg/mL were calculated from the fluorescence intensities by using data extraction analyzer tool (Raybiotech).

### Flowcytometry of bone marrow cells

Mononuclear cells (MNCs) were isolated from the bone marrow by using Ficoll (Cytivia) as described earlier.(46) Cells were prepared for flow cytometry by resuspending in the cell staining buffer with Trustain (Biolegend). Monocytes (CD45^+^Ly6G^-^Ly6C^+^CD115^+^) and macrophages, classically activated (CD45^+^CD11b^+^Ly6G^-^Ly6C^+^F4/80^+^CCR2^+^CD80^+^) or alternatively activated (CD45^+^CD11b^+^Ly6G^-^Ly6C^+^F4/80^+^CX3CR1^+^CD206^+^), were characterized by using the gating strategies as indicated in the Supplementary Figures S1 and S2. Dead cells were excluded by using Aquablue (Invitrogen). Sources and concentrations of different antibodies used in this study are listed in supplementary Table S2.

### Clonogenic Assay

BM-MNCs were cultured in Methocult GF M 3534 medium (StemCell Technologies) according to the manufacturer’s instructions. After 10 days of culture, the colonies were imaged by using Leica microscope and the cells were dissociated from the colonies. Flow cytometry was carried out as described above for monocyte-macrophages sub-populations.

### Macrophage Polarization

MNCs derived from bone marrow were resuspended in RPMI-1640 media and plated at a density of 2 to 3 million cells per well and after 2 hours, the media was removed, and the adherent cells were incubated with RPMI-1640 with 10% FBS,1% penicillin/streptomycin, 1% l-glutamine and 20 ng/mL recombinant mouse M-CSF. The media is changed every 2 days and at day-8 the macrophages (M0) were treated with (100 ng/mL LPS + 10 ng/mL Interferon gamma (IFN-γ)) or (20 ng/mL IL-4 + 20 ng/mL IL-13) for generating classically activated macrophage (CAMs) or alternatively activated macrophages (AAMs), respectively. After two days, the cells were gently harvested and characterized by flow cytometry as described above.

### Treatment of BM-MNCs with cecal supernatants

Cecum from mice were collected in the liquid nitrogen and crushed to fine powder. Cecal content (25 mg) was resuspended in 10 mL of cold DMEM and kept on rocking platform at 200 rpm for 1 hour in cold room (2 - 8°C). The suspension was centrifuged at 3000g for 10 minutes and the cecal supernatant (CS) was filtered by using 0.45 µM filter and then twice with 0.22 µM syringe filter in aseptic conditions. BM-MNCs from young mice were incubated with cecal supernatants derived from different experimental groups at 1:10 dilution for 1 hour and then, mixed with Methocult for CFU-GM assay as described above. Cecal supernatants were also evaluated in the macrophage polarization assay for which, the M0 macrophages from the young BM cells were treated with cecal supernatants for one hour prior to the polarization.

### Data analysis

Results were expressed as Mean ± SD. ‘n’ indicates the number of mice used per each treatment group unless indicated otherwise. Different experimental groups were compared for significant difference by using nonparametric One-way ANOVA, Kruskall-Wallis test followed by Dunn’s test or Two-way ANOVA followed by Bonferroni’s test by using GraphPad Prism software (9.4.0). Treatment groups were considered significantly different if *P* < 0.05.

## Results

Protein expression of MasR and MrgD in the colon tissue-homogenates showed changes with aging. MasR expression was increased in the aging colons in both male and females (*P* < 0.05, n = 6) (Figure 1A and 1B). MrgD expression was decreased in the colons with aging in male mice (*P* < 0.05, n = 6) but not significantly changed in female mice (Figure 1A and 1B). Then, we tested the effect of Ala on the intestinal permeability in the Old mice by using a FITC-dextran permeability assay. A dose-response study indicated that Ala at 0.01 or 0.1 μg/Kg/min perfusion rate for four weeks was not effective in reversing the increased permeability in the aging mice. On the other hand, at the perfusion rate of 1 μg/Kg/min for four weeks, Ala reversed the increased permeability (*P* < 0.0001, n = 4) (Figure 1C). Additional studies were performed with Ala (1 μg/Kg/min, four weeks) in two separate cohorts of male and female mice to confirm the protective effect of Ala in both sexes. Ala treatment reversed the permeability in both male and female Old mice (*P* < 0.0001, n = 7) (Figure 1D and 1E). Concurrent administration of D-Pro^7^-Ang-(1–7) blocked the beneficial effect of Ala (Figure 1D and 1E). It is noteworthy that the increase in the permeability was higher in the Old-female mice than that observed in the Old-males (*P* < 0.05). The reversal of age-associated increase in the intestinal permeability by Ala was further supported by the circulating lipopolysaccharide (LPS)-binding protein (LBP) levels, which were increased in the Old groups compared to the respective Young (*P* < 0.01, n = 7), and were decreased by Ala treatment (*P* < 0.05 (males) or < 0.01 (females), n = 7) (Figure 1F).

**Figure 1.**
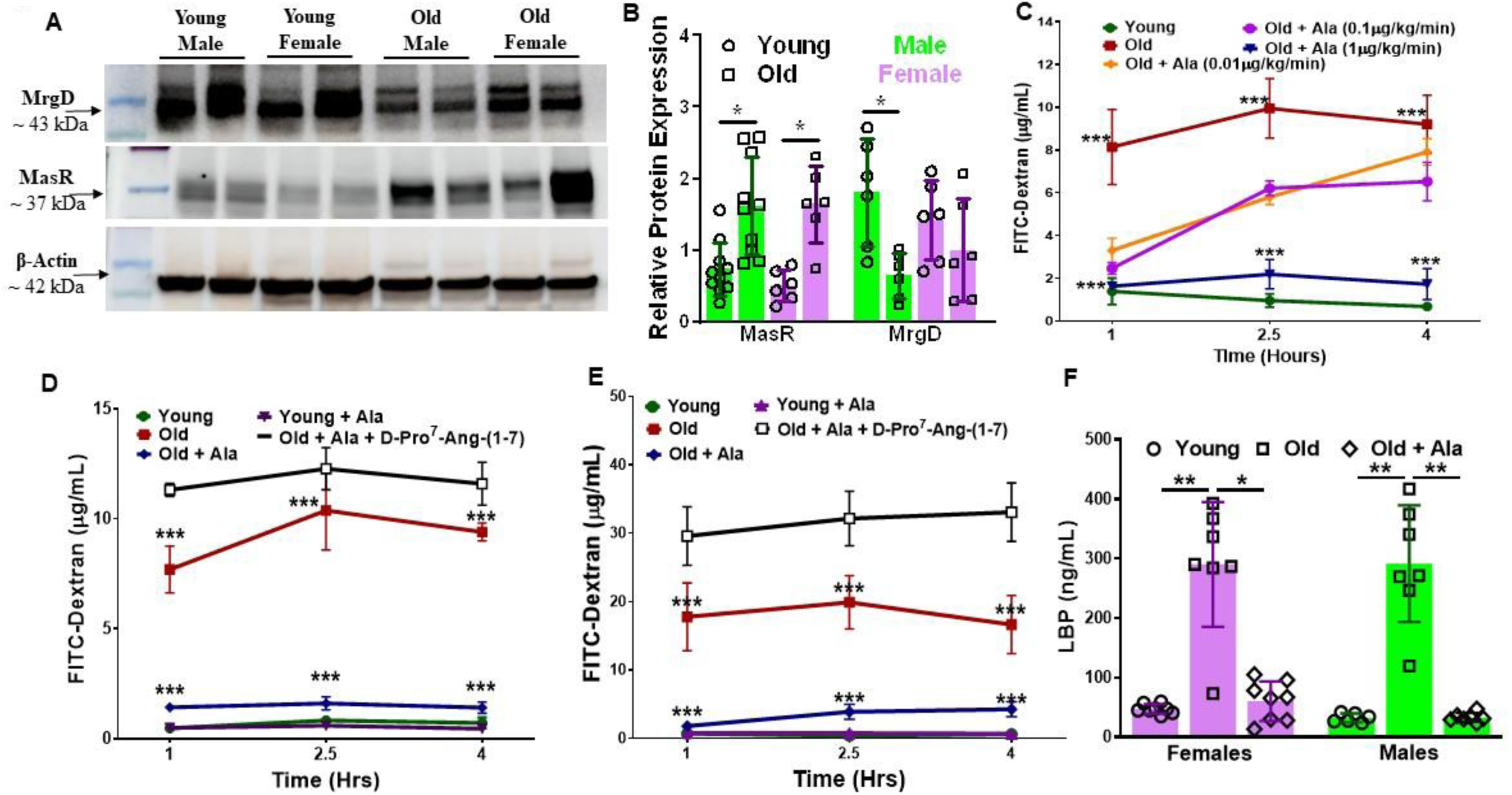
Reversal of the increased intestinal permeability in aging by Alamandine. **A** and **B.** Shown were representative western blots of MrgD, MasR and β-actin proteins in the colon tissue lysates from young and old mice. The expression of MasR was increased in the Old colons of both males and females (*P* < 0.05, n = 6) while that of MrgD was decreased in the male (\**P* < 0.05, n = 6) but unchanged in females compared to the Young. **C**. Dose-dependent effect of Alamandine (Ala) on the intestinal permeability in the Old mice tested by FITC-dextran permeability. Permeability was higher at all three time points tested in the Old group compared to the Young (\*\*\**P* < 0.001, n = 4). At the dose of 1 μg/Kg/min for four weeks, Ala reversed the increased permeability in the Old mice (*P* < 0.001, n = 4) compared to the untreated group. The effect at lower doses, 0.01 and 0.1 μg/Kg/min for four weeks, was not significant (n = 4). **D** and **E**. Ala treatment reversed the increased permeability in both male and female Old mice (\*\*\**P* < 0.001, n = 7). Effect of Ala in the Old group was not observed when concurrently administered with D-Pro^7^-Ang-(1-7) (n = 5). Two-way ANOVA followed by Bonferroni’s multiple comparison was used. **F**. The plasma levels of LBP were higher in the Old (*P* < 0.01, n = 7) compared to the Young. Ala-treatment decreased LBP levels in both male (*P* < 0.05) and females (*P* < 0.01, n = 7) compared to the respective untreated group. Kruskal-Wallis test followed by Dunn’s test was used for statistical comparison.

Histological evaluation revealed shorter and disrupted villi structures in the Old group. Ala-treatment restored the villus structure (Figure 2A). Along similar lines, PAS staining showed decreased number of goblet cells in the villi of Old mice, expressed as numbers per crypt (*P* < 0.05, n = 5) (Figure 2B – 2C), which was associated with decreased mucus thickness (*P* < 0.001, n = 5) (Figure 2B - 2D). Ala-treatment increased the number of goblet cells (*P* < 0.05, n = 5) (Figure 2B - 2D), which accompanied increased thickness of mucus layer (*P* < 0.01, n = 5) (Figure 2B - 2D) (Figure 2B – 2D) in the Old group.

**Figure 2.**
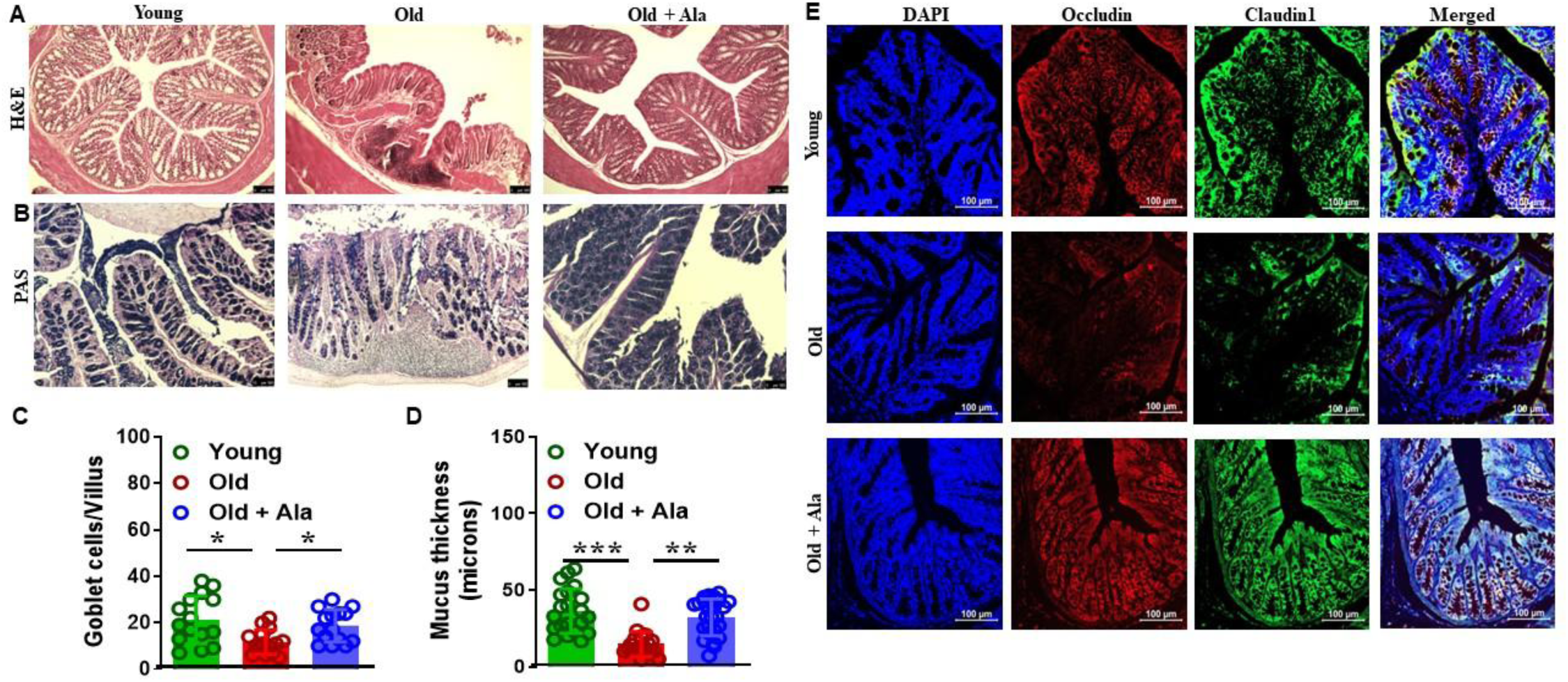
Alamandine restored epithelial barrier integrity in the aging colon. **A**. Shown were representative light microscopic images of Hematoxylin–Eosin (H&E) staining of colon sections. Histological changes depicted disruption of epithelial continuity with atrophied crypts in the Old mice, which was reversed by Ala treatment (n = 5). **B** - **D**. Shown were representative light microscopic images of Alcian Blue PAS staining of colon sections. The number of Goblet cells per villus was decreased in the Old group compared to the Young (\**P* < 0.05, n = 15 villi/5 mice) that was increased by Ala (* *P* < 0.05, n = 15 villi/5 mice). Mucus layer was either thinner or absent in the Old colons compared to the Young (*** *P* < 0.001, n = 6) that was increased by Ala (\*\**P* < 0.001, n = 6). Data sets were analyzed by Kruskall–Wallis test followed by Dunn’s test. **E**. Shown were representative fluorescence images of claudin 1 and occludin immunohistochemistry with nuclear counter stain DAPI in colon sections derived from the experimental groups, Young, Old and Ala treated Old mice. Decreased expression of claudin 1 and occludin in the colons of Old mice were reversed by Ala (n = 6 mice).

Immunofluorescent staining for the tight junction proteins, occludin and claudin 1, showed decreased expression in the Old colons compared to the Young, and was restored by Ala-treatment (Figure 2E).

Then, we examined if the disrupted structural and barrier properties of colon epithelial layer involved alterations in the ISC layer. In the colon sections, MrgD was found to be localized with Lgr5 in the crypts, a marker for ISCs (Figure 3A). Olfm4 and Lgr5 dual-positive cells, ISCs, were found to be decreased in numbers in the Old colons compared to the Young (Figure 3B). Ala-treatment restored the ISC layer in the Old colons (Figure 3B). Then, the expression of MrgD was confirmed in the cultured colon organoids and was colocalized with Lgr5 (Figure 3C). Organoids derived from colons of Old mice were smaller (76 ± 42 microns (min 16 and max 177), *P* < 0.0001, n = 42 organoids/6 mice) compared to those derived from the Young (272 ± 150 microns (min 80 and max 552), n = 36 organoids/6 mice) (Figure 3D and 3E). The colon organoids derived from Ala-treated Old group measured larger (243 ± 118 microns (min 126 and max 533 microns), *P* < 0.0001, n = 39 organoids/6 mice) compared to that derived from the untreated group (Figure 3D and 3E). In agreement with these findings, protein expression of Wnt3a, a critical regulator of ISC homeostasis, was decreased in the Old colons compared to the Young (*P* < 0.01, n = 6), which was increased by Ala-treatment (*P* < 0.01, n = 6 mice/group) (Figure 3F and 3G).

**Figure 3.**
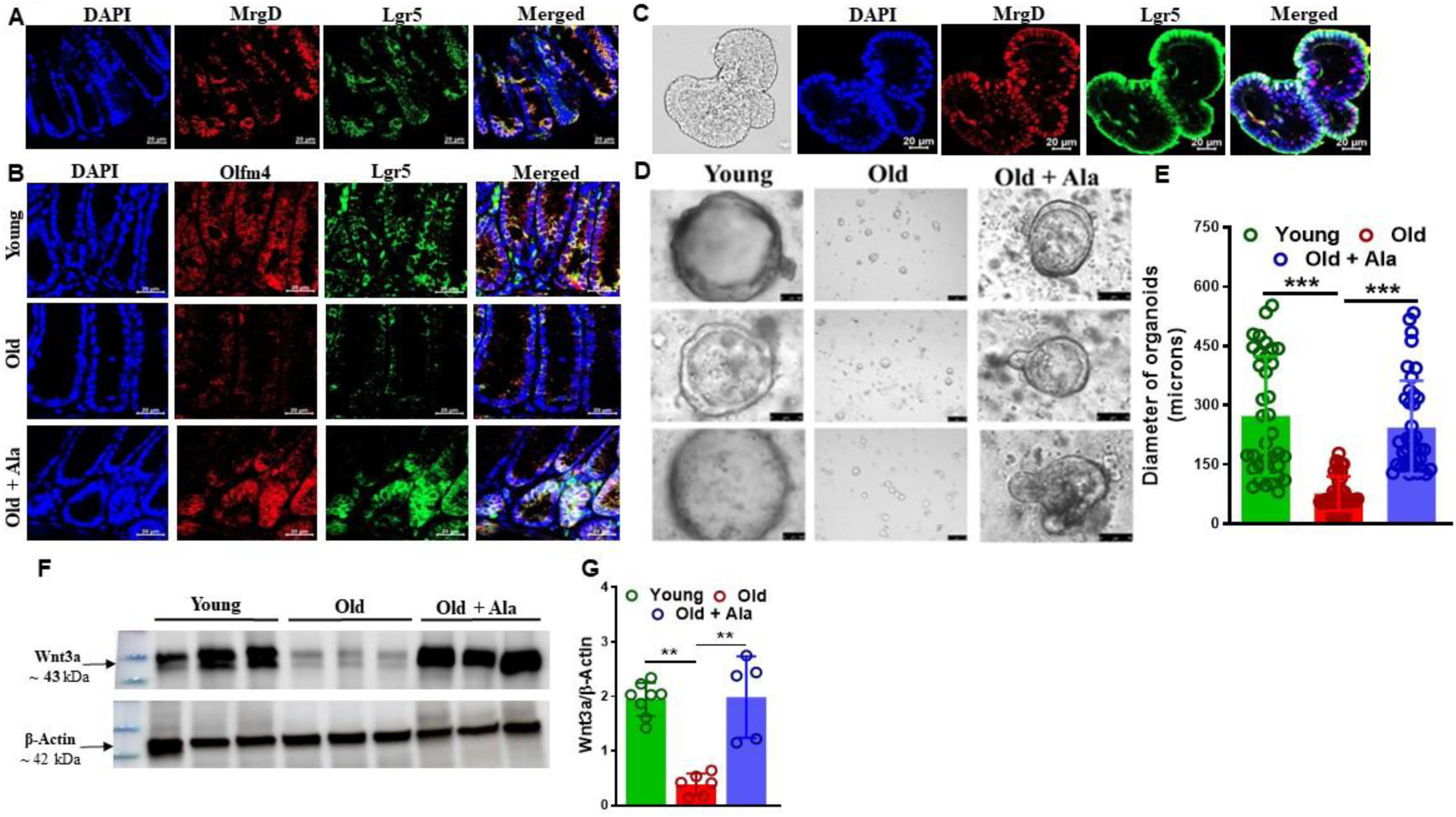
Alamandine restored intestinal stem cell layer in the aging colon. **A**. Shown were representative fluorescence images of MrgD and Lgr5 immunohistochemistry with nuclear counter stain DAPI in colon sections of Young mice showing evidence for the colocalization of MrgD with Lgr5. **B**. Shown were representative fluorescence images of Lgr5 and Olfm4 with nuclear counter stain DAPI in colon sections derived from the experimental groups, Young, Old and Alamandine (Ala) treated Old mice. Decreased Olfm4 and Lgr5 dual positive structures in the Old group was restored by Ala-treatment. **C**. Shown were representative light microscopic image and fluorescent images of colon organoid derive from Young mice with MrgD and Lgr5 immunohistochemistry showing evidence for colocalization of MrgD with Lgr5. **D** and **E**. Shown were representative light microscopic images of colon organoids derived from the experimental groups, Young, Old and Ala treated Old mice. The number and the diameter of organoids were lower in the Old group (\*\*\**P* < 0.001) that were increased by Ala-treatment (\*\**P* < 0.001). **F**. Shown were representative western blots of Wnt3a in the colon lysates derived from three experimental groups, Young, Old and Ala-treated Old mice. Wnt3a expression was decreased in the Old colons (\**P* < 0.01, n = 6 vs Young n = 8) that was restored by Ala-treatment (\*\**P* < 0.01).

In organoids derived from Old-colons, Ala increased Wnt3a in a concentration-dependent manner with similar effects at 10 and 100 nM (Figure 4A and 4C) while Ang-(1–7) did not restore Wnt3a at 100 nM. The changes in the Wnt3a levels induced by the peptides were in agreement with changes observed in claudin1 expression (Figure 4A and 4D). The effect of Ala was significantly antagonized by D-Pro^7^-Ang-(1–7) (*P* < 0.05, n = 5) but partially antagonized by A779, a selective antagonist of MasR that was not significant (Figure 4B and 4C). It is noteworthy that neither of the peptides showed an effect on the Wnt3a levels in the organoids from Young colons (data not shown). Restoration of Wnt3a by Ala in the Old group (*P* < 0.05, n = 10 vs untreated Old) was decreased or reversed by pre-treatment of colon-organoids with a pharmacological inhibitor of GαS (NF449, 10 µM) (*P* < 0.01, n = 5), adenylyl cyclase (SQ22536, 300 µM) (*P* < 0.01, n = 7) or CREB (666-15, 1 µM) (*P* < 0.05, n = 5) (Figure 4E and 4F). None of the blockers produced any changes in the Wnt3a levels, when treated without Ala (Figure 4E and 4F). Phospho-CREB levels were lower in the Old colon-organoids (*P* < 0.001, n = 5) (Figure 4G and 4I). Ala time-dependently activated CREB via phosphorylation in the Old colon-organoids with maximal response at 3-hour treatment (*P* < 0.01, n = 5) (Figure 4G and 4I). Activation of CREB by Ala was completely reversed by D-Pro^7^-Ang-(1–7) (*P* < 0.05, n = 5) but partially antagonized by A779 (Figure 4H and 4J). Along similar lines, active form of β-catenin was increased by Ala in the Old colon-organoids that was blocked by D-Pro^7^-Ang-(1–7) (*P* < 0.05, n = 5) (Figure S3).

**Figure 4.**
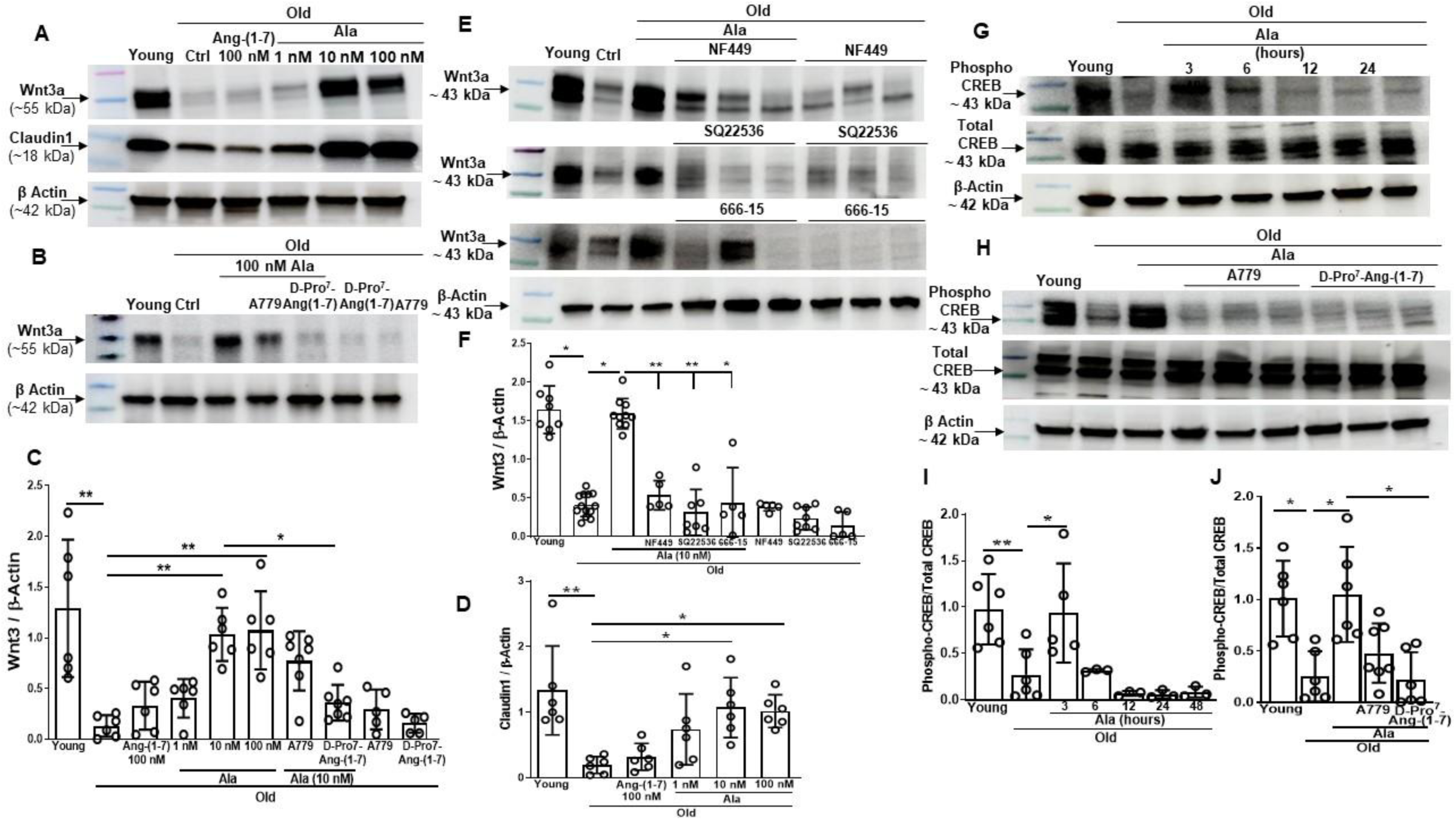
Alamandine restored Wnt3a levels in the colonic organoids derived from Old mice via MrgD/Gαs/AC/CREB/β-catenin pathway. **A** and **B.** Shown were representative western blots of Wnt3a or claudin proteins with β-actin as an internal control in the lysates of colon organoids treated with different concentrations of Ang-(1-7) or Ala with or without A779 or D-Pro^7^-Ang-(1-7). **C - D**. The organoids derived from the Old mice showed lower levels of Wnt3a and claudin1 compared to that of Young mice (\*\**P* < 0.01). Ang-(1-7) (100 nM) and Ala (1 nM) increased Wnt3a and claudin1 that did not achieve statistical significance. Increase in the Wnt3a or claudin1 by Ala at 10 and 100 nM was significantly higher than the untreated Old-colon organoids (\*\**P* < 0.01) and similar to that observed in the Young. Effect of Ala (10 nM) on Wnt3a in the Old-colon organoids was not affected by the simultaneous treatment with A779 (1 µM) but decreased by D-Pro^7^-Ang-(1-7) (1 µM) (\**P* < 0.05). A779 or D-Pro^7^-Ang-(1-7) (1 µM) did not show any increase in the Wnt3a levels of Old-colon organoids. **E**. Shown were representative western blots of Wnt3a and and β-actin proteins in the colon organoids derived from the Young or Old mice, treated with different pharmacological inhibitors. **F**. The organoids derived from the Old mice showed lower levels of Wnt3a and claudin1 compared to that of Young mice (\*\**P* < 0.05, n = 10). Increase in the Wnt3a levels in the Old-colon organoids by Ala was decreased by NF449 (10 µM, \*\**P* < 0.01, n = 5), SQ22536 (300 µM, \*\**P* < 0.01, n = 7) and 666-15 (1 µM, \**P* < 0.05, n = 5). All three blockers showed no change in the Wnt3a levels in the absence of Ala. **G** and **I**. Shown were representative western blots depicting the time-course of CREB phosphorylation by Ala in the colon organoids derived from the Young or Old mice, along with β-actin. Phospho-CREB levels were lower in the Old-colon organoids compared to the Young (\**P* < 0.01, n = 5), which was increased by Ala (10 nM) (\**P* < 0.05, n = 5), after three hours of treatment. **H** and **J**. Shown were representative western blots of Phospho-CREB in the colon organoids treated with Ala with or without antagonists, along with β-actin. Phospho-CREB levels were lower in the Old-colon organoids compared to the Young (\**P* < 0.05, n = 5) that were increased by Ala (10 nM) (\**P* < 0.05, n = 5). The effect of Ala was inhibited by D-Pro^7^-Ang-(1-7) (1 µM) (\**P* < 0.05) but not by A779 (1 µM) (n = 5).

Then, we tested if the restoration of epithelial barrier integrity was able to rescue aging-associated gut microbial dysbiosis. The richness of gut microbiome represented by Chao1 was decreased in the Old mice, as reflected by Chao1 index compared with Young (*P* < 0.01, n = 7) and was restored by Ala treatment (*P* < 0.05, n = 7) (Figure 5A). Beta diversity visualized by PCoA using weighted Bray-Curtis distances demonstrated distinct clustering of samples from three groups (*P* = 0.0001, F: 4.633, One-way PERMANOVA). While Young and Old groups clustered distantly (*P* = 0.0003, n = 7), Ala-treated Old group clustered closer to the Young group (*P* = 0.0009) (Figure 5B) as detected by the multiple comparisons. Spearman’s rank correlations with Benjamini-Hochberg correction was used to identify correlations between microbial families and functional end points of gut barrier integrity. FITC-Dextran permeability was positively correlated with the abundance of *Peptostreptococcaceae, Closstridaceae and Deferribacteraceae* while negative correlation was observed with *Erysipelotrichaceae*, *Lactobacillaceae*, *Bacteroidaceae*, *Tannerellaceae*, *Peptococcaceae* and *Acholeplasmataceae*. LPS levels were positively correlated with the abundance of *Clostridaceae* while negatively correlated with *Lactobacillaceae*, *Bacteroidaceae*, *Tannerellaceae*, *Peptococcaceae*, *Eubacteriaceae* and *Monoglobaceae* (Figure 5C).

**Figure 5.**
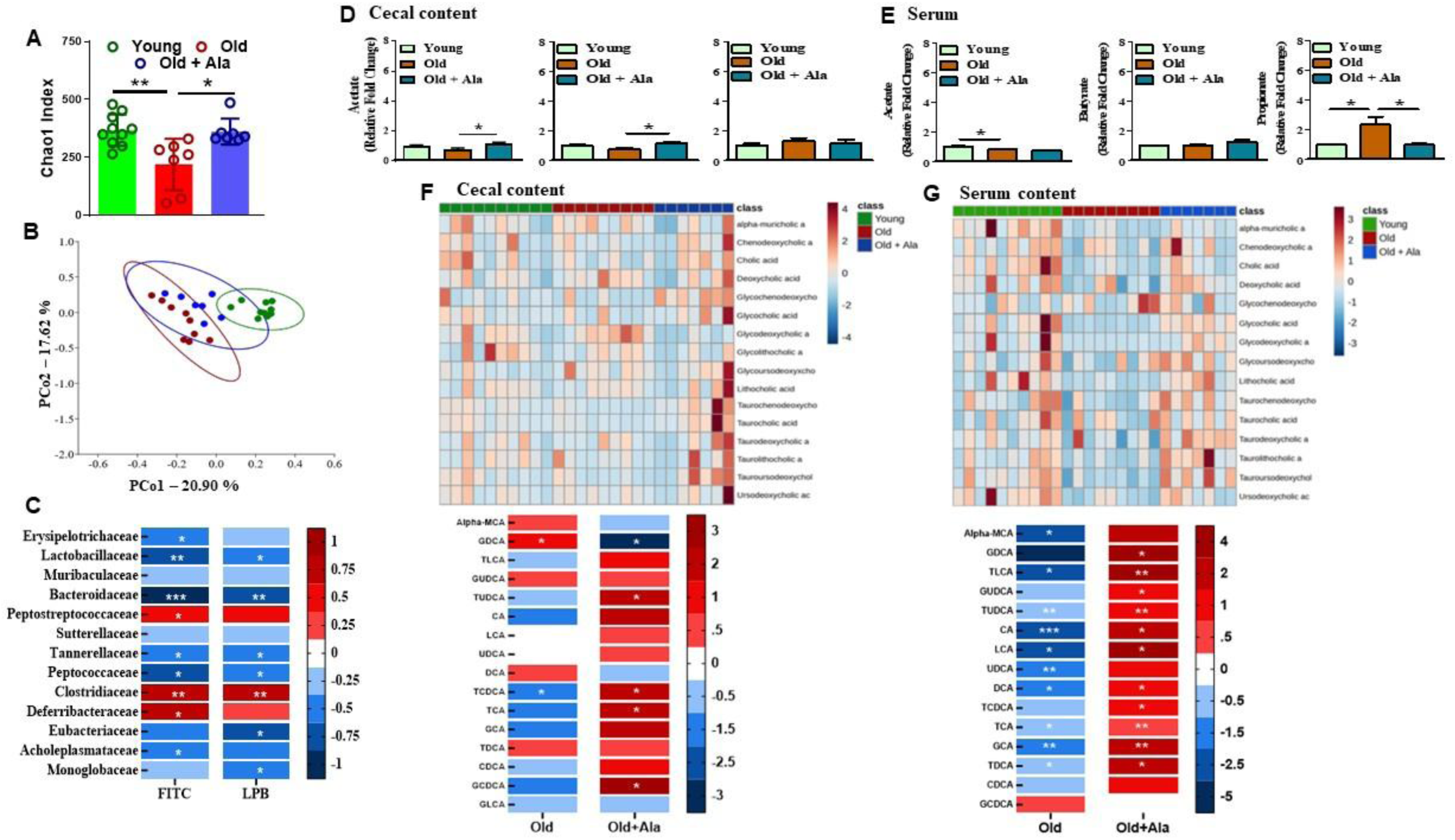
Restructuring of dysbiotic gut microbiota in the aging by Alamandine. A. Histograms showing a decrease in the richness (Chao1 index) of the microbiota in the Old group compared with the Young, which was restored by Ala. B. Beta diversity as tested by principal coordinate analysis by using Brey–Curtis was significantly different among three groups depicting differences in microbial composition (PERMANOVA analysis). **C**. Correlation of microbial families with functional measures of gut permeability carried out by Spearman’s rank correlations (\**P* < 0.05, \*\**P* < 0.01 and \*\*\**P* < 0.001). **D** and **E**. Gut microbial metabolites, acetate and butyrate were higher in cecum of Old mice treated with Ala (\**P* < 0.05) compared with the untreated. In the serum, the propionate levels were higher in the Old compared to the Young that was decreased by Ala-treatment. Kruskal-Wallis test followed by Dunn’s test. **F.** Shown were heat map and comparison of the concentrations of bile acid metabolites in the cecum. The levels of GDCA and TCDCA were higher or lower, respectively, in the Old groups compared to the Young (\**P* < 0.05, n = 10). The levels of GDCA was decreased and that of TUDCA, TCDCA, TCA and GCDCA were increased (\**P* < 0.05, n = 7) with Ala treatment in the cecal content of Old mice compared to the untreated group. **G**. Shown were heat map and comparison of the concentrations of bile acid metabolites in the serum. The metabolites, Alpha-MCA, TLCA, TUDCA, CA, LCA, UDCA, DCA, TCA, GCA and TDCA, that were decreased in the Old group compared to the Young (n = 10), were increased by Ala-treatment (n = 7) with the exception of UDCA. While GDCA, GUDCA and TCDCA were unchanged in the Old compared to the Young but increased by Ala. \**P* < 0.05 or \*\**P* < 0.01, Kruskal-Wallis test followed by Dunn’s test.

Serial ANOVA tests were carried out to detect differences at the taxonomic level of phylum, family and genus. At the phylum level (Figure S4), relative abundance of *Bacillota* was increased in the Old compared with Young and decreased by Ala treatment but neither of these changes achieved statistical significance. In contrast, the relative abundance (RA) of *Bacteroidota* was decreased in the Old compared to the Young group (*P* < 0.05) and was increased by Ala (*P* < 0.01). Gut microbial dysbiosis in the Old group characterized by increased *Bacillota/Bacteroidota,* which was reversed by Ala (*P* < 0.05). The RA of *Pseudomonadota* was increased in the Old group compared to the Young and was reversed by Ala treatment. Along similar lines, the increased RA of *Verrucomicrobiota* in the Old (*P* < 0.01 vs Young) was reversed by Ala (*P* < 0.05 vs untreated Old).

At the family level (Figure S4), the RA of *Lactobacillaceae, Muribaculaceae* and *Bacteroidaceae* was decreased in the Old compared to the Young, which was increased by Ala (*P* < 0.05 - 0.01). In contrast, the RA of *Erysipelotrichaceae* was decreased in the Old (*P* < 0.05 vs Young) but was not altered by Ala. The RA of *Rikenellaceae* showed increased trend with aging, which was further increased by Ala (*P* < 0.05 vs untreated Old) while the increased RA of *Clostridaceae* (*P* < 0.01) was reversed by Ala (*P* < 0.01 vs untreated Old).

At the genus level (Figure S4), the RA of *Lactobacillus* was decreased in the Old (*P* < 0.001 vs Young) and was restored by Ala (*P* < 0.05). *Muribaculum* showed decreased trend in the Old compared to the Young and was further decreased by Ala (*P* < 0.05). *Bacteroides* was not detected in the Old group, but Ala treatment restored with an RA of 1.4±0.3%. In aged mice, *Clostridium* was found to be higher compared to the Young (*P* < 0.01 vs Young) and was decreased by Ala (*P* < 0.01). The genus *Dubosiella* was decreased in the Old (*P* < 0.001 vs Young) but was not altered by Ala. Alternatively, an unresolved genus within *Ruminococcaceae* was decreased in the Old compared to Young (*P* < 0.05) and was increased by Ala (*P* < 0.05 vs untreated old). Lastly, Ala increased the RA of *Butyricicoccus* (*P* < 0.05 vs untreated Old) that was decreased in the Old compared to the Young.

Then, we tested the cecal and the circulating bacterial metabolites. Among the short chain fatty Acids (SCFAs) tested, acetate and butyrate were increased in the cecum (*P* < 0.05, n=7), which were not observed in the circulation. Propionate levels were unchanged in the cecum but were increased in the circulation of the Old compared to the Young (*P* < 0.05, n=7), which was reversed by Ala (*P* < 0.05, n=7) (Figure 5D and 5E). Along similar lines, several changes were observed in the bile acid levels in the cecum and the circulation. Glycodeoxycholic acid (GDCA) was increased in the Old cecal supernatants, which was reversed by Ala while it was increased by Ala in the circulation (Figure 5F and G). Tauroursodeoxycholic acid (TUDCA), Taurochenodeoxycholic acid (TCDCA) and Taurocholic acid (TCA) were either unchanged or decreased in the Old ceca and were increased by Ala in cecal supernatants as well as in the circulation (Figure 5F and G). Glycochenodeoxycholic acid (GCDCA) was increased by Ala in the ceca but unchanged in the circulation. Taurolithocholic acid (TLCA), Taurodeoxycholic acid (TDCA), Glycoursodeoxyxcholic acid (GUDCA), cholic acid (CA), Glycocholic acid (GCA), Lithocholic acid (LCA), Ursodeoxycholic acid (UDCA), Deoxycholic acid (DCA) and TCDCA, and alpha-muricholic acid (alpha-MCA) were similar in ceca from all three groups while decreased or unchanged in the circulation of the Old group, which was reversed by Ala (Figure 5F and G). Chenodeoxycholic acid (CDCA) levels were unchanged in either the cecal supernatants or in the circulation in all experimental groups (Figure 5F and G). Glycolithocholic acid (GLCA) levels were unchanged in the cecum among three groups and was not detected in the serum.

Colon inflammation was evaluated by immunohistochemical examination of monocyte-macrophage infiltration in the colon sections and by analyzing the inflammatory factors in the tissue lysates. Pro-inflammatory CD80^+^ macrophages were higher (*P* < 0.01, n = 5) while anti-inflammatory CX3CR1^+^ macrophages were lower (*P* < 0.01, n = 5) in the colon sections of Old mice compared to the Young (Figure 6A-6C). These changes were reversed by treatment with Ala (Figure 6A-6C). The following inflammatory factors, IL-13, MCP-5, CXCL1, CXCL4, CXCL9, CXCL13, MIP-1a, CCL17 and Eotoxin 2 (CCL24) were all higher in the colon lysates from the Old group (*P* < 0.05 to 0.001) although CXCL1 was not significant (Figure 6D). All of these changes were reversed by Ala treatment (*P* < 0.05) although the effect on CCL24 and CCL17 was not statistically significant (Figure 6D). Similar trends were observed in the Old group compared to the Young or in the Old upon Ala-treatment in the levels of IL17, IL15, CCL5, TNFα, Eotoxin (CCL11), CD30L, MIP1 and TIMP1 but did not achieve significance while no changes were noted in the levels of IL10, IL15, IL21, TNFRII, LIX and MCP1 (Figure 6D).

**Figure 6.**
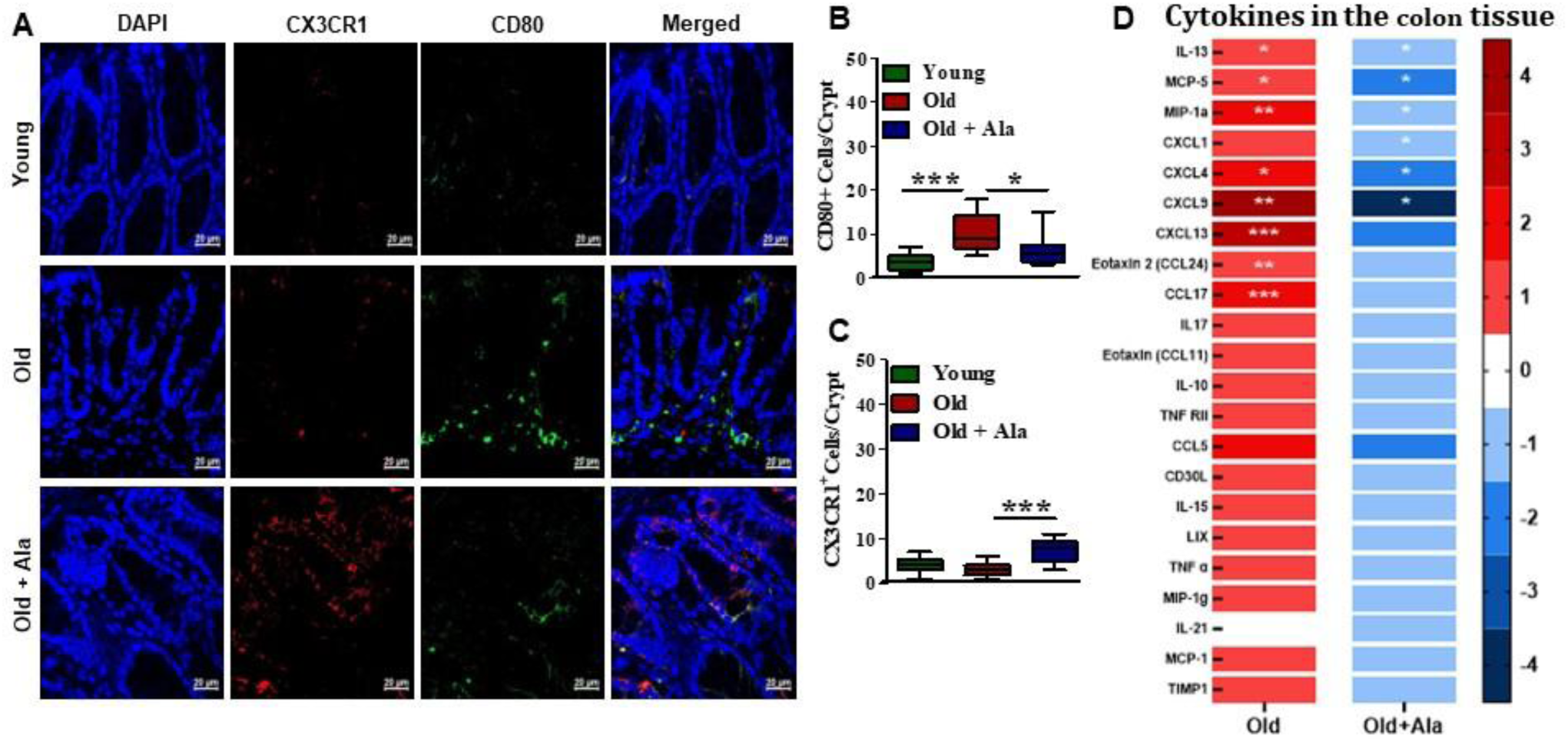
Aging-associated gut wall inflammation is ameliorated by Alamandine. **A.** Shown were representative fluorescence images of CD80 and CX3CR1 immunohistochemistry with nuclear counter stain DAPI in colon sections from different treatment groups. **B** and **C**. Increased number of CD80 cells in the old mice colon were decreased while the number of CX3CR1 cells were increased by Ala treatment in old mice (\**P* < 0.05, \*\**P* < 0.001 vs old untreated, n = 6). **D**. The tissue concentrations of cytokines such as IL-13, MCP-5, MIP-1a, CXCL4 CXCL9, CXCL13, CCL24 and CCL17 were higher in colons from the Old mice compared to the Young, which were decreased by Ala. Effect of Ala on CXCL13, CCL24 and CCL17 was not significant. CXCL1 was higher in the Old colons compared to the Young but was not significant and it was significantly decreased by Ala. \**P* < 0.05, \*\**P* < 0.001, Kruskal-Wallis test followed by Dunn’s test was used for statistical significance.

In agreement with the previous studies, aging bone marrow was characterized by pro-inflammatory environment with higher number of Ly6G^-^CD11b^+^Ly6C^+^ monocytes (*P* < 0.001, n=7) and M1 (CD80^+^CCR2^+^) Mϕs (*P* < 0.001), and lower number of anti-inflammatory M2 (CX3CR1^+^CD206^+^) Mϕs (*P* < 0.001) compared to the Young with an increased M1/M2 ratio (*P* < 0.01) (Figure 7A-7E). MrgD is expressed in the BM-stem/progenitor cells and no change was observed with age (Figure S3). The observed changes in the monocyte-macrophage populations in the Old group were reversed by treatment with Ala except in the M1 Mϕs resulting in the decreased ratio of M1/M2 Mϕs (Figure 7A-7E). Similar findings were observed in the female mice (Figure S6).

**Figure 7.**
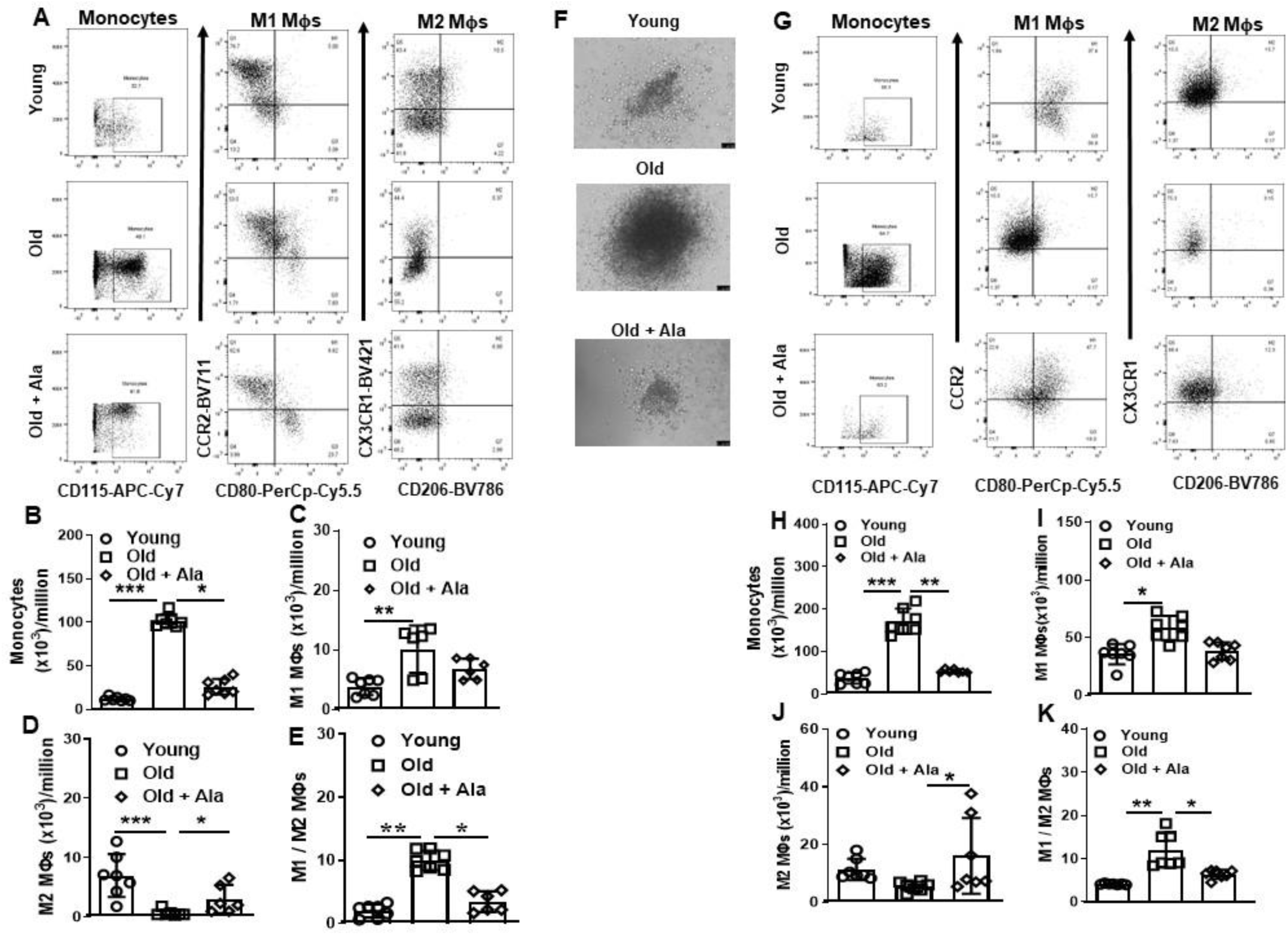
Aging-associated increase in the myelopoiesis in the bone marrow was decreased by Alamandine. **A**. Shown were representative flow cytometry dot plots of CD45^+^Ly6G^-^CD11b^+^Ly6C^+^CD115^+^ monocytes, CD45^+^Ly6G^-^CD11b^+^Ly6C^+^F4/80^+^CD80^+^CCR2^+^ Mϕ (M1) and CD45^+^Ly6G^-^CD11b^+^Ly6C^+^F4/80^+^CD206^+^CX3CR1^+^ Mϕ (M2) in the bone marrow from three experimental groups of mice. **B - E**. Monocytes and M1 Mϕs were higher in the Old group compared to the Young (\*\*\**P* < 0.001 and \*\**P* < 0.01, respectively) that were decreased by Ala-treatment (\**P* < 0.05) though the decrease was not significant in M1 Mϕs. M2 Mϕs were decreased in the Old group (\*\*\**P* < 0.001) that resulted in the higher ratio of M1/M2 Mϕs. Ala-treatment increased the number of M2 Mϕs and decreased the ratio (\**P* < 0.05). **F**. Shown were representative light microscopy images of colonies derived from the bone marrow cells undergoing CFU-GM assay. **G**. Shown were representative flow cytometry dot plots of monocytes, M1 Mϕ and M2 Mϕ in the colonies derived from three experimental groups of mice. **H** – **K**. Monocytes and M1 Mϕs were higher in the colonies derived from Old mice compared to the Young (\*\*\**P* < 0.001 and \**P* < 0.05, respectively). Monocytes were decreased (\*\**P* < 0.01) but M1 Mϕs were not significantly affected by Ala-treatment. M2 Mϕs were decreased in the Old group that was increased by Ala and these changes were not significant. A higher ratio of M1/M2 Mϕs was observed in the Old group compared to the Young (\*\**P* < 0.01) that was decreased by Ala-treatment (\**P* < 0.05).

*In vitro* myelopoiesis by CFU-GM assay showed that the colonies derived from the BM progenitor cells of the Old group were larger than that observed in the colonies from Young BM (Figure 7F). Colonies derived from the Ala-treated Old mice were of similar size to that observed in the Young (Figure 7F). Flow cytometry of the dissociated cells from the colonies revealed higher number of monocytes (*P* < 0.001, n=7), M1 Mϕs (*P* < 0.05, n=7) from the Old group and a lower number of M2 Mϕs (not statistically significant) compared to those obtained from the Young resulting in an increased ratio of M1/M2 Mϕs (*P* < 0.001) (Figure 7G and 7K). Ala treatment decreased the monocytes (*P* < 0.01) and increased the M2 Mϕs (*P* < 0.05) with no change in the M1 Mϕs and yet resulting in a decreased ratio of M1/M2 Mϕs (*P* < 0.05) (Figure 7G and 7K). Similar findings were observed in the female mice (Figure S6).

Lastly, the effect of cecal supernatants (CS) from three groups on the myelopoietic potential of BM-stem/progenitor cells was tested. Treatment of BM cells from Young mice with CS derived from Old mice (O-CS) and subjected to CFU-GM assay, resulted in higher number of monocytes (*P* < 0.05, n = 6-7) and lower number of M2 (CX3CR1^+^CD206^+^) (*P* < 0.05, n = 6-8) Mϕs but no significant change in the pro-inflammatory M1 (CD80^+^CCR2^+^) Mϕs compared to that treated with CS derived from Young mice (Y-CS) (Figure 8A – 8E). The ratio of M1/M2 was higher with O-CS (*P* < 0.05, n=6-8) (Figure 8F). The CS derived from Ala-treated Old mice (O-Ala-CS) decreased the monocytes (*P* < 0.01, n=7) and M1 Mϕs (*P* < 0.01, n=6-7) but no change in the M2 macrophages, which resulted in the lower ratio of M1/M2 Mϕs (*P* < 0.01, n=6-8) (Figure 8A – 8F). Similar findings were observed in the macrophage polarization assay on the induction of pro- and anti-inflammatory Mϕ phenotypes. O-CS induced higher number of M1 Mϕs (*P* < 0.01, n = 5) and lower number of M2 (*P* < 0.05, n = 5) Mϕs compared to that induced by Y-CS while O-Ala-CS had no effects on M1 Mϕ but increased the M2 Mϕs (*P* < 0.05, n = 5) (Figure 8G – 8J) that resulted in a lower ratio of M1/M2 Mϕs.

**Figure 8.**
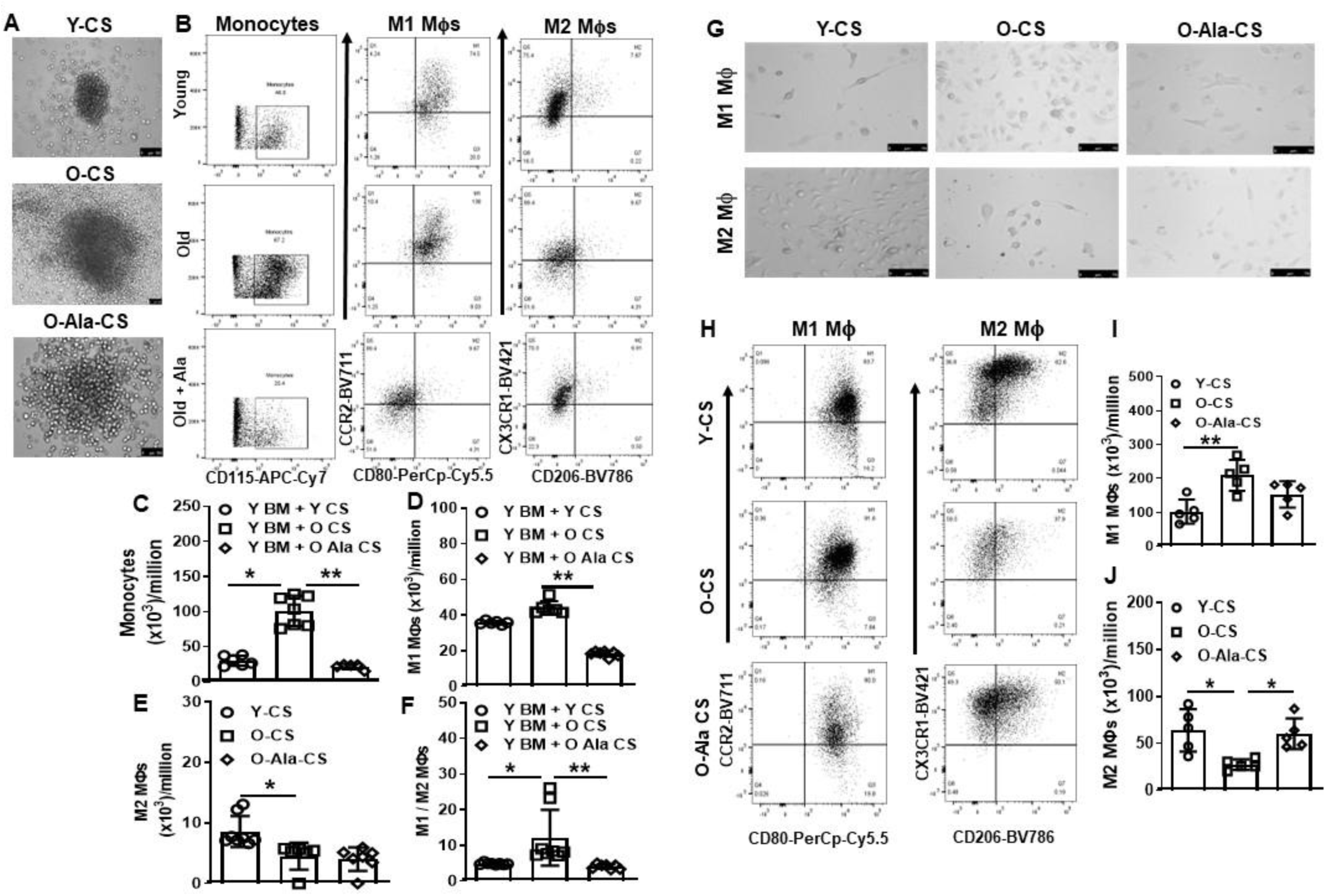
Pro-myelopoietic potential of bacterial metabolites in the aging was reversed by Alamandine. **A**. Shown were representative light microscopy images of colonies derived from the bone marrow cells treated with cecal supernatants (CS) from three different experimental groups and subjected to CFU-GM assay. **B**. Shown were representative flow cytometry dot plots of CD45^+^Ly6G^-^CD11b^+^Ly6C^+^CD115^+^ monocytes, CD45^+^Ly6G^-^CD11b^+^Ly6C^+^F4/80^+^CD80^+^CCR2^+^ Mϕ (M1) and CD45^+^Ly6G^-^CD11b^+^Ly6C^+^F4/80^+^CD206^+^CX3CR1^+^ Mϕ (M2) derived from CFU-GM colonies. **C** – **F**. Monocytes were higher in the Old-CS (O-CS) treated group compared to the Young-CS (Y-CS) (\**P* < 0.05) that were lower in the Old-Ala-CS (O-Ala-CS) treated group (\*\**P* < 0.01). M1 Mϕs showed increasing trend in the O-CS group but was not significant. M2 Mϕs were decreased in the O-CS group (\**P* < 0.05) that resulted in the higher ratio of M1/M2 Mϕs (\**P* < 0.05). O-Ala-CS treatment did not alter the number of M2 Mϕs but decreased the ratio (\**P* < 0.05). **G**. Shown were representative light microscopy images of Mϕs derived from the bone marrow cells treated with cecal supernatants (CS) from three different experimental groups and subjected to Mϕ polarization assay. **H**. Shown were representative flow cytometry dot plots of monocytes, M1 and M2 Mϕs (M2) derived from BM cells treated with CS from different experimental groups and subjected to Mϕ polarization assay. **I** and **J**. M1 Mϕs were higher in the O-CS treated group compared to the Y-CS (\*\**P* < 0.01) that were lower in the group treated with O-Ala-CS, which did not achieve significance. M2 Mϕs were lower in the O-CS treated group compared to the Y-CS (\**P* < 0.05) that were increased in the O-Ala-CS group (\**P* < 0.05).

## Discussion

This study reports several novel findings. MrgD is expressed in the ISCs located in the crypts of colonic mucosa as well as in the stem/progenitor cells of mice and the expression is decreased in the aging colon. Ala regenerated ISCs in the aging colon and restored epithelial barrier integrity by acting on MrgD. Ala activated Wnt3a/β-catenin signaling in the organoids from aging colon. MrgD is coupled to Gαs/Adenyl cyclase/cAMP/PKA/CREB pathway to induce Wnt3a in the colon organoids in aging. Ala-treatment restructured dysbiotic gut microbiota in aging and altered cecal and circulating levels of gut microbial metabolites. Ala-treatment ameliorated inflammation in the colon and increased the number of anti-inflammatory macrophages in the bone marrow and colon. Importantly, Ala conferred anti-inflammatory phenotype to the microbial metabolomic profile. Collectively, these results provide compelling evidence for the pharmacological potential of Ala/MrgD pathway to restore homeostasis in the gut-bone marrow axis in aging.

In our study MrgD expression is decreased in the Old colon while MasR expression was increased none the less the receptor expression was still present in the aging colons. Increased MasR expression is in agreement with our previous reports in aging or diabetic colons,(10) which we described as a compensatory upregulation of the receptor expression for the protective angiotensin, Ang-(1–7). Studies in other tissues showed either an increase or a decrease in the MrgD protein expression. Decreased expression was observed in the hearts following transverse aortic constriction.(47) However, downregulation was reported in the pulmonary arterial smooth muscle cells (PASMCs) derived from mouse models of pulmonary hypertension and in PASMCs derived from human subjects with PAH yet beneficial effects of Ala were evident.(24) In agreement with this, in the current study, despite the lower levels of MrgD expression, Ala dose-dependently decreased the intestinal permeability in the Old mice *in vivo*, which was further supported by decreased systemic levels of LBP, a reliable marker of intestinal epithelial barrier integrity. Complete reversal of this effect of Ala by D-Pro7-Ang-(1–7), an antagonist of MrgD,(19) confirmed that the effect is mediated by MrgD.

In agreement with the previous report (8), Wnt3a levels and ISCs were decreased in the aging colons of male mice. In the female mice however despite relatively higher permeability than that in males, Wnt3a levels showed a decreased trend that was not significant. Further investigation is needed to delineate the involvement of other isoforms of Wnt in the observed sex-difference. Increased Wnt3a expression paralleled the expression of claudin1, one of several targets of Wnt3a via increasing active β-catenin levels. Ang-(1–7) was found to be less potent at increasing the Wnt3a levels in the Old colons despite the upregulation of MasR expression. We have previously shown that Ang-(1–7) restored gut barrier integrity and restored Wnt3a levels in the aging colons. *In vitro* studies in the current study indicated that Ang-(1–7) is relatively less potent than Ala in restoring Wnt3a in the Old colon-organoids. It is likely that Ang-(1–7) is converted to Ala in the gut or in the systemic circulation to Ala prior to acting on the gut mucosa however we do not have evidence in support of this. While ACE2 was shown to generate Ala from Ang A,(19, 20) the likely enzyme that can convert Ala to Ang-(1–7) was hypothesized to be aspartate β-decarboxylase, which is yet to be investigated. It is noteworthy to mention that a hypothesis was put forth for the involvement of microbial origin of a yet to be identified enzyme with decarboxylase activity that can metabolize Ang-(1–7) to Ala. Increased Wnt3a levels by Ala were blocked by D-Pro7-Ang-

(1–7) but not by A779, a selective antagonist of MasR.(48, 49) It is important to note that D-Pro^7^-Ang-(1–7) was first reported as an antagonist at MasR (50) and that Ala and Ang-(1–7) were shown to act via two different receptors.(51) Furthermore, in another study the effect of Ala was competed by β-alanine and was antagonized by D-Pro7-Ang-(1–7).(19) Alamandine produced endothelium-dependent vasorelaxation that was blocked by D-Pro^7^-Ang-(1–7), but not by A779.(19) In agreement with this, Tetzner et al (25) showed that alamandine activated cAMP formation in endothelial and mesangial cells transfected with MrgD. In the current assay system using colon organoids and pharmacological antagonists, the effect of Ala appears to be mediated via acting on MrgD but not MasR however further investigation is required to support this conclusion.

Effect of Ala on Wnt3a is further confirmed by the concurrent increase in the claudin expression and active β-catenin levels. Overall increased ISC activity as reflected by the fully developed organoids from Old colons contributed to the restoration of different cell layers of colon wall, including epithelial and mucosal layers and importantly, the barrier integrity. Active β-catenin mediates the transcriptional response to the interaction of Wnt with the cognizant receptor, frizzled receptor. In the absence of Wnt3, β-catenin is phosphorylated at Ser^33^/Ser^37^or/Thr^41^ by glycogen synthase kinase-3 (GSK-3) followed by ubiquitination and degradation.(5, 52) Elevated intracellular Wnt levels stabilizes cytoplasmic β-Catenin in the active form, which upon translocation to the nucleus induce the Wnt-target gene expression that stimulate regeneration of ISCs and expression of Lgr5, claudin1, occludin and several others. (7, 52, 53)

MrgD was shown to mediate cellular signaling events via multiple G-proteins including Gi, Gs and Gq-proteins, depending on the cell type. However, in this study we focused at Gαs-dependent signaling as the expression of Wnt proteins is known to be regulated by cAMP/protein kinase A (PKA)/CREB pathway (54, 55). Our studies using pharmacological inhibitors of Gαs, adenylyl cyclase and CREB, NF449, SQ22536 and 666-15, respectively, confirmed the involvement of Gαs/AC/PKA/CREB pathway in the expression of Wnt3a by Ala/MrgD interaction. This is further confirmed by increased levels of active CREB in organoids derived from Ala-treated Old colons. In agreement with the above studies, the increased active CREB levels were blocked by D-Pro^7^-Ang-(1–7) not by A779.

The increased permeability in the aging mouse colons was associated with higher infiltration of pro-inflammatory macrophages indicated by CD80^+^ positivity and decreased anti-inflammatory CX3CR1^+^ cells, which were reversed by Ala-treatment. The inflamed colon in the aging was associated with upregulation of nine different pro-inflammatory factors, which may directly contribute to the increased permeability and inflammation-induced downregulation of ISC functions.(56) Earlier studies showed evidence for anti-inflammatory mechanism of Ala in ameliorating vascular remodeling that was associated with decreased pro-inflammatory (IL1β, TNF and CCL2) and pro-fibrotic factors (MMP2 and TGF β1), and increased pro-resolution markers (MRC1 and FIZZ1) (57).

Amelioration of inflammation and the restoration of the barrier integrity induced remarkable changes in the aging gut microbiome, which collectively resulted in the reversal effects on inflammaging. The anti-inflammatory effect on the aging colon was correlated with the effects of Ala observed in the BM-myelopoiesis and Mϕ polarization towards anti-inflammatory phenotype. Mϕs are heterogeneous population of cells largely consisting of classically activated (M1) Mϕs (high IL-12 and low IL-10) which are induced by IFNγ and microbial products and alternatively activated (M2) Mϕs (low IL-12 and high IL-10) induced by IL-4 and IL-13.(58, 59) The current and a previous study (60) detected the MrgD protein in the mouse stem/progenitor cells and human macrophages, respectively. This study reports a reversal of increased myelopoiesis in aging mice and restoration of balance in the dysregulated macrophage polarization in both males and females. The study by Rukavina Mikusic et al (60) showed evidence for the anti-inflammatory function of Ala/MrgD pathway in LPS-stimulated human macrophages by decreasing the secretion of IL-6 and IL-β1. Consistent with these effects of Ala, changes were observed in the local and systemic pro-inflammatory profile. BM in the aging mice demonstrated increased myelopoiesis resulting in the higher number of circulating monocytes and importantly, skewing of macrophage polarization towards pro-inflammatory phenotype. All of these pro-inflammatory changes were reversed by Ala.

The levels of microbe-associated molecular patterns (MAMPs), such as LPS, were found to be elevated in the circulation of aging mice. (61, 62) This study evaluated LBP, an acute-phase protein that interacts with LPS and transfer monomers to the receptor complex on immune cells and induce the production of pro-inflammatory factors.(63, 64) LBP is a stable indicator of LPS exposure.(65, 66) Our study showed increased levels of LBP in the aging mice were decreased by Ala most likely by restoring the colon epithelial barrier integrity, which in turn contributed to the decreased colon inflammation. Ala increased the beta-diversity and decreased the F/B ratio in the Old group suggesting restructuring of the microbiome. *Bacillota*/*Bacteroidota* predicts the circulating levels of TMAO and the TMA-generating bacteria.(67) TMAO promotes myelopoiesis and macrophage polarization toward pro-inflammatory phenotype that elevates local and systemic inflammatory milieu.(17, 68) The decreased TMAO in the circulation and increased *Bacteroidota* infers that the effects of Ala on myelopoiesis and macrophage polarization are likely due to the restructured gut microbiota.

Our study confirmed that the bacterial metabolites as enriched in the cecal supernatants collectively induce myelopoiesis and pro-inflammatory macrophages. Our study detected significant changes in the microbiota-derived metabolites that are known to modulate gut inflammation via modifying Mϕ polarization. Selected short-chain fatty acids and bile acids were analyzed in the cecum as well as in the serum. Cecal concentration of metabolites did not correlate with serum concentrations, which can be attributed to influences by metabolism, kinetics and enterohepatic circulation. Among the three SCFAs, only propionate was increased in the Old group that was reversed by Ala. A study by Wang et al (69) showed that propionate can activate NLRP inflammasome in the presence of microbiota-derived TLR-ligands in macrophages and stimulated IL-1β release.

Earlier studies showed strong evidence for the modulation of Mϕ-polarization by BAs largely acting on TGR5. Free and conjugated BAs are produced by the liver enzymes from cholesterol and the bacterial enzymes, bile salt hydrolase and 7-dehydrolase deconjugate BAs and convert primary BAs to secondary BAs, respectively.(70, 71) BAs are now known to function as signaling molecules via acting on receptors, farnesoid X receptor (FXR) and Takeda G protein receptor 5 (TGR5). (72, 73) The latter mediates the actions of BAs on macrophages and myelopoietic cells. (74) Among the three experimental groups we find significant differences in thirteen of the sixteen bile acids that were analyzed. Bile acids, DCA and LCA, were shown to activate TGR5 in monocyte-Mϕs that counter-regulates CD14/TLR4 pathway and decrease the release of pro-inflammatory factors TNFα, IL1β and IL6.(75) LCA, TLCA and TDCA decreased M1 pro-inflammatory macrophages while increasing the M2 macrophages.(76, 77) While we are not aware of studies that support the effects of individual metabolites on macrophages, we believe that the observed effects of Ala resulted from the collective effect of all the factors that were found to be increased or decreased. It is important to note that Ala/MrgD activation was shown to modulate macrophage polarization and pro-inflammatory potential in vitro, (60) which was independent of microbial metabolites therefore the direct actions of Ala on monocyte-macrophages via MrgD cannot be ruled out.

This study however was not free from limitations. Ala was administered by using osmotic pumps which is not always clinically feasible however an approach involving micellar formation by using β-cyclodextrin for oral administration was previously demonstrated in rodents. Study did not involve quantitative analysis of Ala in the circulation or in the tissues. Feasibility of the Ala quantification in biological fluids by HPLC was reported therefore further investigation is required to determine the pharmacokinetic properties of Ala.

In conclusion, this study provides strong evidence for the pharmacological potential of Ala/MrgD pathway in restoring the colon barrier integrity via activating the regenerative functions of ISCs as well as by restoring the homeostasis of gut-BM axis.

## List of abbreviations used in the text

ACE: Angiotensin-converting enzyme
Ala: Alamandine
Ang-(1-7): Angiotensin-(1-7)
AT1R: Angiotensin II type 1 receptor
APC: Allophycocyanin
BM: Bone marrow
BCA: Bicinchoninic acid
BSA: Boven serum albumin
CX3CR1: C-X3-C motif chemokine receptor 1
CXCL1: CXC motif chemokine ligand 1
CFU-GM: Colony-forming unit – granulocyte and macrophages
CS: Cecal supernatant
DAPI: 4’,6-diamidino-2-phenylindole
Fas-L: Fas ligand
FITC: Fluorescein isothiocyanate
GSK-3: Glycogen synthase kinase-3
HSC: Hematopoietic stem cells
IFNγ: Interferon-gamma
IL: Interleukin
ISC: Intestinal stem cells
LBP: lipopolysaccharide-binding protein
Lgr: Leucine-rich repeat containing G protein-coupled receptor
LPS: Lipopolysaccharide
MasR: Mas receptor
MCP-1: Monocyte chemoattractant protein-1
M-CSF: Macrophage colony-stimulating factor
MD2: Myeloid differentiation protein 2
MIP1α: Macrophage inflammatory protein 1 alpha
MNC: Mononuclear cells
MrgD: Mas related G-protein coupled receptor, member D
Olfm: Olfactomedin
PBS: Phosphate buffered saline
PE: Phycoerythrin
PVDF: Polyvinylidene fluoride
RA: Relative abundance
RAS: Renin angiotensin system
SDS/PAGE: Sodium dodecyl-sulfate/polyacrylamide gel electrophoresis
TARC: Thymus and activation-regulated chemokine
TLR4: Toll-like receptor 4
TMAO: Trimethylamine N-oxide
TNFα: Tumor necrosis factor alpha
Wnt3a: Wingless and Integration 1 proteins, member 3a

## Data availability

The 16S rRNA amplicon sequencing data reported in the current manuscript are freely available at the National Center for Biotechnology Information (NCBI) Sequence Read Archive (SRA) as BioProject PRJNA1347378.

Other data sets supporting the results reported in this article will be available upon request from the authors.

## Author contributions: CRediT statement

KC Methodology and Data curation

ZLM, VM and SA Methodology

HER Data curation

AAE Resources and Project administration

YPRJ Conceptualization, Investigation, Resources, Project administration, Supervision and Writing.

