## Supplement Tables and Figures for "Alamandine/MrgD Pathway Modulates Gut-Bone Marrow Axis in Aging"

**Running title**: Alamandine/MrgD and the leaky gut of aging

*__Address for correspondence__:

Yagna PR Jarajapu, M Pharm, Ph D, FAHA.
Sudro-16, Albrecht Blvd.,
Department of Pharmaceutical Sciences,
College of Health and Human Sciences,
North Dakota State University,
Fargo, North Dakota, USA.


**Table S1. List of antibodies used for IHC and western blotting studies.**

| Antibody | Catalog # | Vendor | Concentration |
|---|---|---|---|
| **Claudin 1 (Host - Ms)** | 37-4900 | Thermofisher | 1:100 |
| **Occludin (Host -Rb)** | 40-4700 | Thermofisher | 1:100 |
| **Lgr5 (Host – Ms)** | MA5-25644 | Thermofisher | 1:100 |
| **Olfm4 (Host – Rb)** | PA5-115687 | Thermofisher | 1:50 |
| **633 Goat Anti Rb** | 20122 | Biotium | 1:250 |
| **633 Goat Anti Ms** | 20010 | Biotium | 1:250 |
| **488 Goat Anti Rat** | A11006 | Invitrogen | 1:100 |
| **DAPI** | 40043 | Biotium | - |
| **MAS1 (G-1)** | Sc- 390453 | Santacruz | 1: 1000 |
| **MrgD** | NC2589487 | Thermo fischer | 1: 1000 |
| **MrgE** | TA316024 | Origene | 1: 1000 |
| **Wnt6** | 501734260 | Thermo fischer | 1: 1000 |
| **Wnt3a** | TA801736S | Origene | 1:1000 |

**Table S2: Antibodies used for flow cytometry and immunohistochemistry of monocyte-macrophage populations.**

| Antibody | Catalog # | Vendor | Concentration |
|---|---|---|---|
| **Aqua Blue** | L34966A | Invitrogen | 0.1 µL/100 µL |
| **CD45 – BV605** | 103140 | Biolegend | 0.5 µL/100 µL |
| **Ly6G – PE-CF594** | 562700 | BD Biosciences | 0.125 µL/100 µL |
| **CD11b – APC eFluor 780** | 47-0112-82 | Invitrogen | 0.06 µL/100 µL |
| **Ly6C – FITC** | 553104 | BD Biosciences | 0.1 µL/100 µL |
| **CD115 – APC/Cy 7** | 135532 | Biolegend | 0.2 µL/100 µL |
| **F4/80 – PE** | 123110 | Biolegend | 0.75 µL/100 µL |
| **CX3CR1 – BV421** | 149023 | Biolegend | 0.2 µL/100 µL |
| **CCR2 – BV711** | 747964 | BD Biosciences | 0.725 µL/100 µL |
| **CD80 – PerCP/Cy5.5** | 104722 | Biolegend | 0.5 µL/100 µL |
| **CD206 – BV785** | 141729 | Biolegend | 0.375 µL/100 µL |
| **DAPI** | 40043 | Biotium | - |
| **633 Goat Anti Rb** | 20122 | Biotium | 1:250 |
| **488 Goat Anti Mouse** | 20010 | Biotium | 1:250 |
| **CD80** | 66406-1-IG | Thermofisher | 1:100 |
| **CX3CR1** | 14-6093-81 | Thermofisher | 1:100 |

**Supplementary figures - legends**

**Figure S1**. **Gating strategy for the characterization and enumeration of monocytes by flow cytometry**. Shown were representative flow cytometry dot plots. Top and bottom panels represent characterization of cells with fluorescent-conjugated isotype controls or the antibodies, respectively.

**Figure S2**. **Gating strategy for the characterization and enumeration of macrophages by flow cytometry**. Shown were representative flow cytometry dot plots. Top and bottom panels represent characterization of cells with fluorescent-conjugated isotype controls or the antibodies, respectively.

**Figure S3**. **Alamandine restored Wnt3a levels in the colonic organoids derived from Old mice via MrgD/Gα$_s$/AC/CREB/β-catenin pathway**. **A**. Shown were representative western blots for active β-Catenin and phospho- β-Catenin in the Old-colonic organoids. **B**. Ratio of active β-Catenin to phospho- β-Catenin in the old organoids was significantly lower compared to the Young (* $P < 0.05$, n = 5) and was increased by Ala treatment (*$P < 0.05$ vs untreated Old, n = 7). The effect of Ala was decreased by D-Pro$^7$-Ang-(1-7) (1 µM) that did not achieve significance compared to the Ala + D-Pro$^7$-Ang-(1-7) group.

**Figure S4**. **Restructuring of dysbiotic gut microbiota in aging by Ala.**
See the main text for details. Kruskal-Wallis test followed by Dunn's test was used for statistical comparisons.

**Figure S5**. **MrgD protein expression in the bone marrow stem/progenitor cells.**
**A** and **B**. Shown were representative western blots of MrgD and β-actin proteins in the bone marrow stem/progenitor cells of either Young or Old mice. No change was observed in the expression of MrgD in the bone marrow stem/progenitor cells with aging.

**Figure S6**. **Aging-associated increase in the myelopoiesis was decreased by Alamandine in female mice.** **A**. Shown were representative flow cytometry dot plots of CD45$^+$Ly6G$^-$CD11b$^+$Ly6C$^+$CD115$^+$ monocytes, CD45$^+$Ly6G$^-$CD11b$^+$Ly6C$^+$F4/80$^+$CD80$^+$CCR2$^+$ Mϕ (M1) and CD45$^+$Ly6G$^-$CD11b$^+$Ly6C$^+$F4/80$^+$CD206$^+$CX3CR1$^+$ Mϕ (M2) in the bone marrow from three experimental groups of female mice. **B - E**. Monocytes and M1 Mϕs were higher in the Old group compared to the Young (*$P < 0.05$ and **$P < 0.01$, respectively) that were decreased by Ala-treatment (*$P < 0.05$). M2 Mϕs were decreased in the Old group (*$P < 0.05$) that resulted in the higher ratio of M1/M2 Mϕs (***$P < 0.001$). Ala-treatment showed no effect in the number of M2 Mϕs and yet decreased the ratio of M1/M2 Mϕs (*$P < 0.05$). **F**. Shown were representative light microscopy images of colonies derived from the bone marrow cells undergoing CFU-GM assay.

**G**. Shown were representative flow cytometry dot plots of monocytes and M1 and M2 Mϕs in the colonies from three experimental groups of mice. **H** – **K**. Monocytes and M1 Mϕs were higher in the colonies derived from Old mice compared to the Young (**$P$ < 0.01 and *$P$ < 0.05, respectively). Monocytes showed decreasing trend while M1 Mϕs were significantly decreased by Ala (**$P$ < 0.01). No changes were observed in M2 Mϕs among three groups. A higher ratio of M1/M2 Mϕs was observed in the Old group compared to the Young (**$P$ < 0.01) that was decreased by Ala-treatment (*$P$ < 0.05).

*Supplement*

**Figure S1**

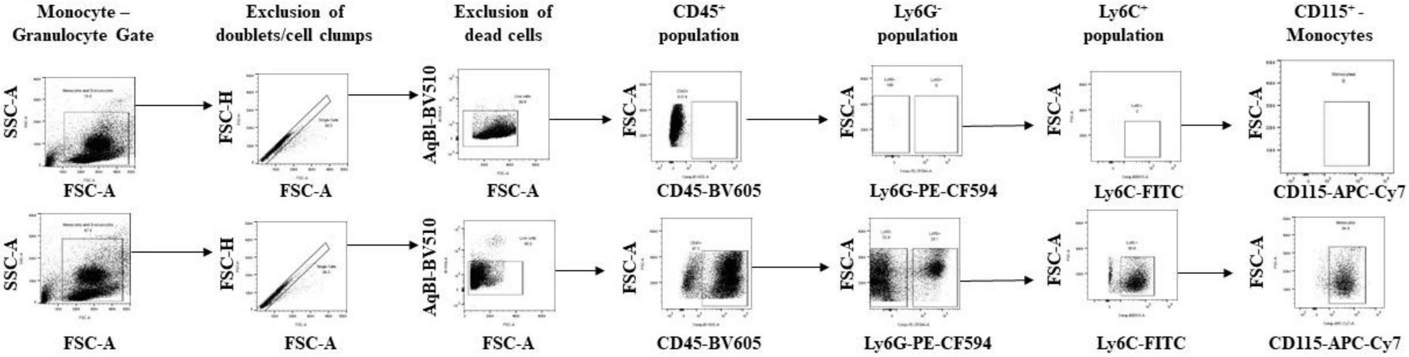

*Supplement*

**Figure S2**

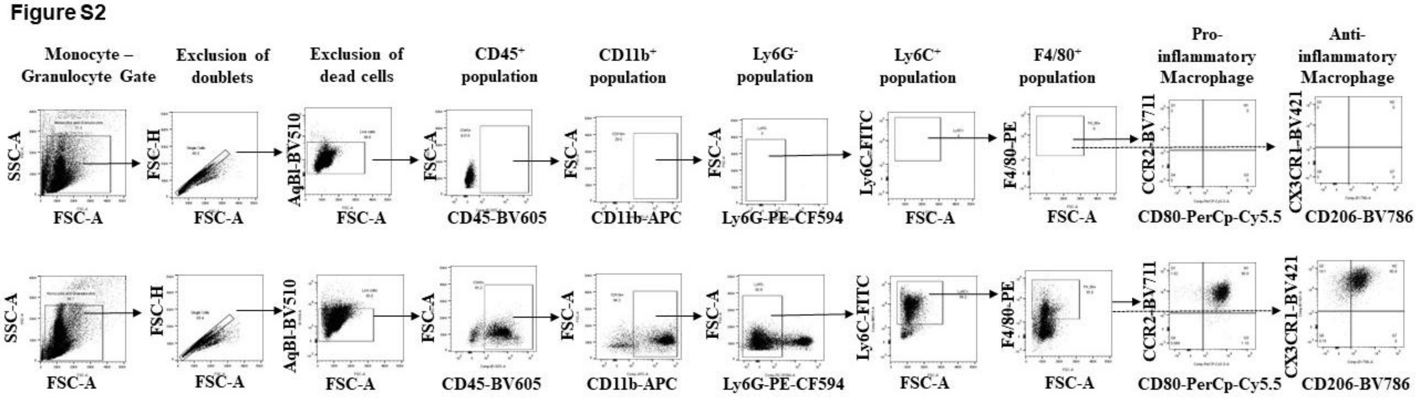

**Figure S3**

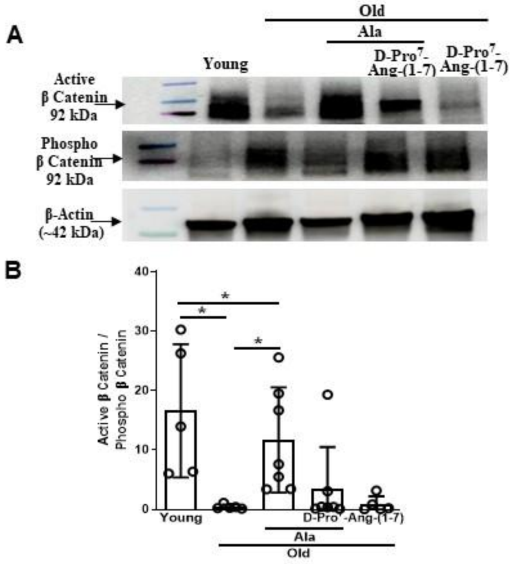
A
Old
Ala
Young
D-Pro7-Ang-(1-7)
D-Pro7-Ang-(1-7)
Active β Catenin 92 kDa
Phospho β Catenin 92 kDa
β-Actin (~42 kDa)
B
Active β Catenin / Phospho β Catenin
40
30
20
10
0
*
*
*
Young
Ala
D-Pro7-Ang-(1-7)
Old

*Supplement*

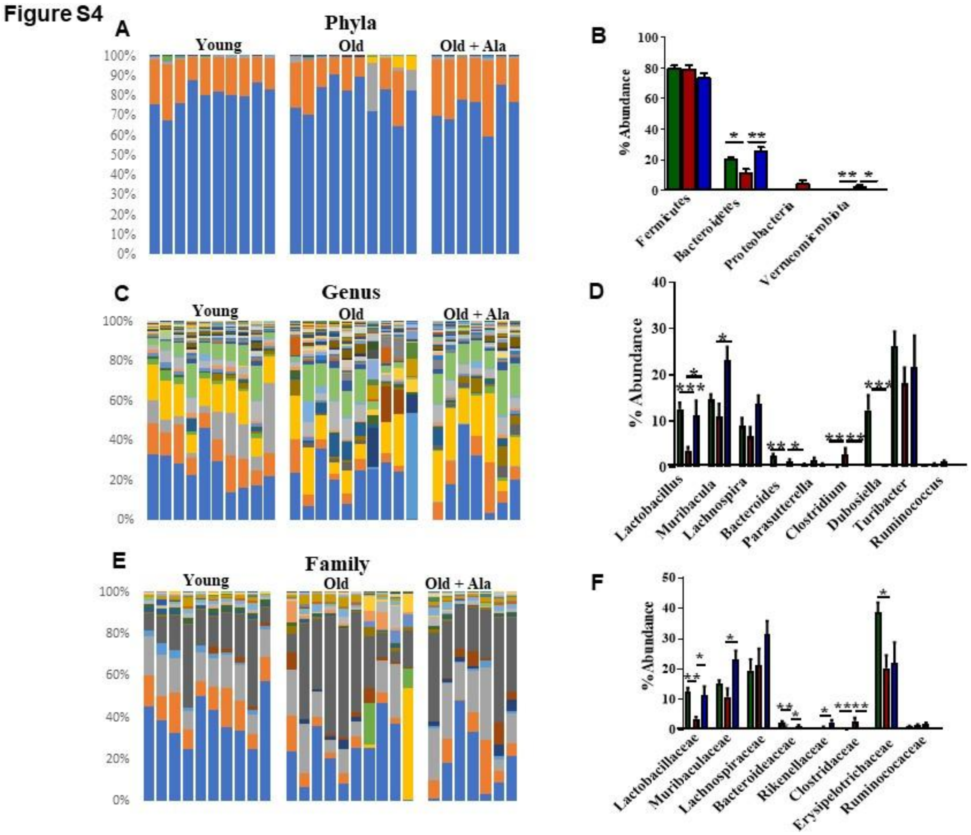
Figure S4
A
Phyla
Young
Old
Old + Ala
B
% Abundance
Firmicutes
Bacteroidetes
Proteobacteria
Verrucomicrobiota
C
Genus
D
Lactobacillus
Muribacula
Lachnospira
Bacteroides
Parasutterella
Clostridium
Dubosiella
Turibacter
Ruminococcus
E
Family
F
Lactobacillaceae
Muribaculaceae
Lachnospiraceae
Bacteroideaceae
Rikenellaceae
Clostridaceae
Erysipelotrichaceae
Ruminococaceae

*Supplement*

**Figure S5**

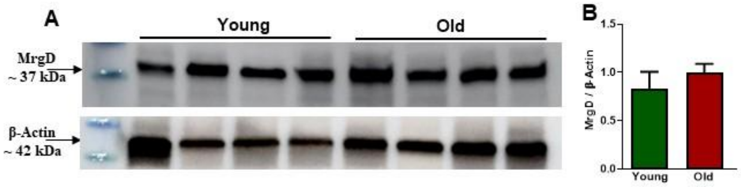

**Figure S6**

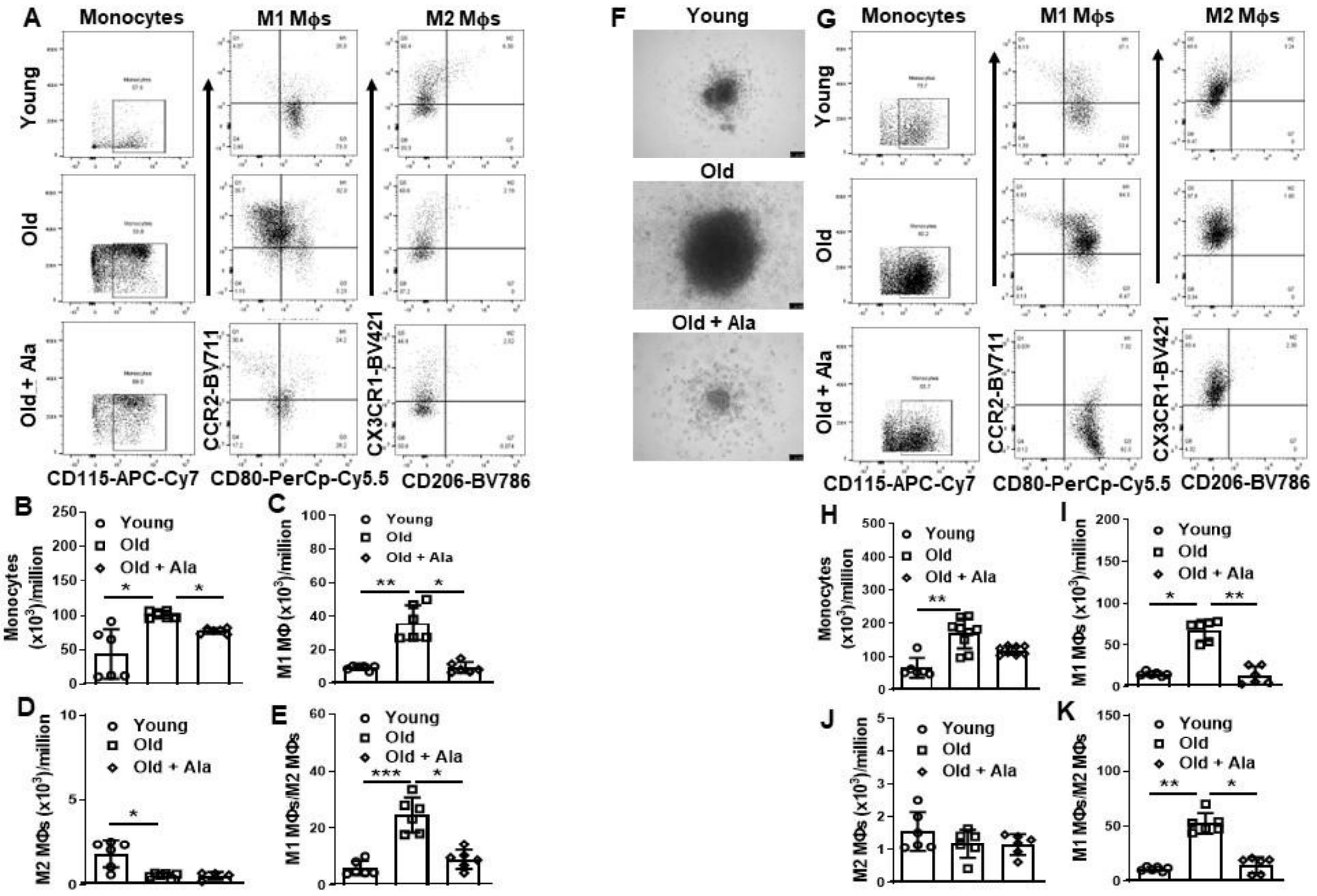
A
Monocytes
M1 Mϕs
M2 Mϕs
Young
Old
Old + Ala
CD115-APC-Cy7
CCR2-BV711
CD80-PerCp-Cy5.5
CX3CR1-BV421
CD206-BV786
F
Young
Old
Old + Ala
G
B
C
D
E
H
I
J
K
Young
Old
Old + Ala
Monocytes (x10$^3$)/million
M1 MΦ (x10$^3$)/million
M2 MΦs (x10$^3$)/million
M1 MΦs/M2 MΦs
M1 MΦs (x10$^3$)/million
